# Mutation of NEKL-4/NEK10 and TTLL genes opposes loss of the CCPP-1 deglutamylase and prevents neuronal ciliary degeneration

**DOI:** 10.1101/2020.05.21.108449

**Authors:** Kade M. Power, Jyothi S. Akella, Amanda Gu, Jonathon D. Walsh, Sebastian Bellotti, Margaret Morash, Winne Zhang, N. Ross, Andy Golden, Harold E. Smith, Maureen M. Barr, Robert O’Hagan

## Abstract

Ciliary microtubules are subject to post-translational modifications that act as a “Tubulin Code” to regulate motor traffic, binding proteins and stability. In humans, loss of CCP1, a <u>c</u>ytosolic <u>c</u>arboxy<u>p</u>eptidase and tubulin deglutamylating enzyme, causes infantile-onset neurodegeneration. In *C. elegans,* mutations in *ccpp-1*, the homolog of CCP1, result in progressive degeneration of neuronal cilia and loss of neuronal function. To identify genes that regulate microtubule glutamylation and ciliary integrity, we performed a forward genetic screen for suppressors of ciliary degeneration in *ccpp-1* mutants. We isolated the *ttll-5(my38)* suppressor, a mutation in the <u>t</u>ubulin <u>t</u>yrosine <u>l</u>igase-<u>l</u>ike glutamylase gene. We show that mutation in *ttll-4, ttll-5,* or *ttll-11* gene suppressed the hyperglutamylation-induced loss of microtubules and kinesin-2 mislocalization in *ccpp-1* cilia. We also identified the *nekl-4(my31)* suppressor, an allele affecting the NIMA (Never in Mitosis A)-related kinase NEKL-4/NEK10. In humans, NEK10 mutation causes bronchiectasis, an airway and mucociliary transport disorder caused by defective motile cilia. *C. elegans* NEKL-4 does not localize to cilia yet plays a role in regulating axonemal microtubule stability. This work defines a pathway in which glutamylation, a component of the Tubulin Code, is written by TTLL-4, TTLL-5, and TTLL-11; is erased by CCPP-1; is read by ciliary kinesins; and its downstream effects are modulated by NEKL-4 activity. Identification of regulators of microtubule glutamylation in diverse cellular contexts is important to the development of effective therapies for disorders characterized by changes in microtubule glutamylation. By identifying *C. elegans* genes important for neuronal and ciliary stability, our work may inform research into human ciliopathies and neurodegenerative diseases.

## Introduction

Cilia and flagella are antenna-like organelles that are conserved from algae to humans and play important roles in sensation, intercellular communication, development, and homeostasis [1]. The core of both cilia and flagella is the axoneme, a conserved microtubule structure with a ring of nine doublets that may surround inner microtubule singlets in motile cilia [1]. The Tubulin Code hypothesis suggests that post-translational modifications, such as glutamylation, acetylation, glycylation, detyrosination, and others, can increase the diversity of microtubules beyond incorporation of different tubulin isotypes for structural and functional specialization [2].

Glutamylation, or addition of side-chains of the amino acid glutamate, of tubulin tails is emerging as an important factor in organizing the microtubule cytoskeleton [3,4]. Microtubules in neurons and cilia can be reversibly glutamylated. Although the function of glutamylation and other post-translational modifications is not completely understood, enzymes that regulate post-translational glutamylation of microtubules are known. Cytosolic carboxypeptidases (CCPs) such as CCP1 deglutamylate microtubules [5,6], opposing the activity of tubulin tyrosine ligase-like (TTLL) enzymes, which add glutamates to tubulin tails [7–9]. Glutamylation side-chains can be as short as a single glutamate, or as long as 20 glutamates [10]. Particular CCP enzymes may reduce the length of glutamate side-chains, and some can remove the branch point glutamate to eliminate glutamylation [5]. Similarly, some TTLL family members initiate glutamate side-chains, while others elongate [9]. TTLLs may also have preferences for α or β tubulin substrates [9]. Therefore, microtubule glutamylation is not uniform, and both side-chain length and tubulin substrate may exert different influences over motors, microtubule-associated proteins (MAPs), or other regulators of microtubule stability.

Glutamylation is essential for neuronal survival and axoneme stability, and a growing list of reports link hyper- or hypo-glutamylation to disease. Dysregulated microtubule glutamylation causes infantile-onset neurodegeneration and the ciliopathy Joubert syndrome in humans, and progressive cerebellar purkinje cell degeneration (*pcd*) in mice [3,5,11–14]. Defects in axonal transport, which may be regulated by the Tubulin Code, contribute to neurodegenerative diseases including Alzheimer’s disease and Parkinson’s disease [4,15]. Although the importance of regulated glutamylation on ciliary and neuronal microtubules are clearly established, a comprehensive understanding of microtubule glutamylation is necessary to understand and interpret its context-specific effects in different cilia and neuron types.

*C. elegans* offers advantages to unraveling the Tubulin Code. Most basic biological processes, including axon development, neuronal transport, and ciliogenesis are conserved between *C. elegans* and humans [16]. In the worm, cilia are located on the distal most ends of sensory dendrites [17–19], are not essential for viability, and are diverse in terms of the composition and arrangement of microtubules, tubulin post-translational modifications, and motor proteins [2,18,20]. In male-specific ciliated sensory neurons, the CCPP-1 deglutamylase and TTLL-11 glutamylase control ciliary ultrastructure and select ciliary kinesin motors [21,22]. In amphid and phasmid sensory neurons, loss of CCPP-1 leads to degeneration of ciliary microtubules indicating cell-specific differences in the tubulin code [21,22].

In *C. elegans* amphid and phasmid neurons, ciliary integrity is easily tested using a dye-filling assay [18]. *ccpp-1* are dye-filling defective (Dyf) in adults but not early larval stages, reflecting ultrastructural defects and progressive ciliary microtubule degeneration [21]. Using a candidate gene approach, we found that deletion mutations in *ttll-4, ttll-5,* or *ttll-11* glutamylase genes suppressed the *ccpp-1* Dyf phenotype. To identify new molecules and pathways that function in ciliary homeostasis, we performed a genetic screen for mutations that suppress the *ccpp-1* Dyf phenotype. We identified mutations in *ttll-5* and in the never-in-mitosis kinase gene *nekl-4*. In humans, TTLL5 mutation causes recessive retinal dystrophy [23,24]. *C. elegans* NEKL-4 is homologous to human NEK10, which is a kinase specific to ciliated cells and contributes to ciliogenesis in cultured human kidney cells [25]. In humans, NEK10 mutation causes bronchiectasis, a disorder of mucociliary transport in the airway due to defective motile cilia [26]. Here we show that NEKL-4 is expressed in all ciliated neurons but does not localize to cilia, suggesting that NEKL-4 indirectly influences regulation of ciliary stability by CCPP-1 and glutamylation. This work represents a step toward elucidating the molecular pathways by which glutamylation, as a component of the Tubulin Code, regulates cilia; this, in turn, allows us to better understand ciliopathies and neuronal survival in humans.

## Results

### Isolation of *ccpp-1* Dyf suppressors

*ccpp-1* mutation causes hyperglutamylation and progressive ciliary degeneration in amphid and phasmid neuronal cilia [21]. Ciliary degeneration is detectable using dye-filling assays, in which *C. elegans* nematodes are soaked in the fluorescent lipophilic dye DiI (**Fig. 1a**). The dye is only taken up in neurons that have intact sensory cilia; therefore, adult *ccpp-1* do not take up this fluorescent dye and display the Dye-filling defective, or “Dyf,” phenotype [21].

**Figure 1.**
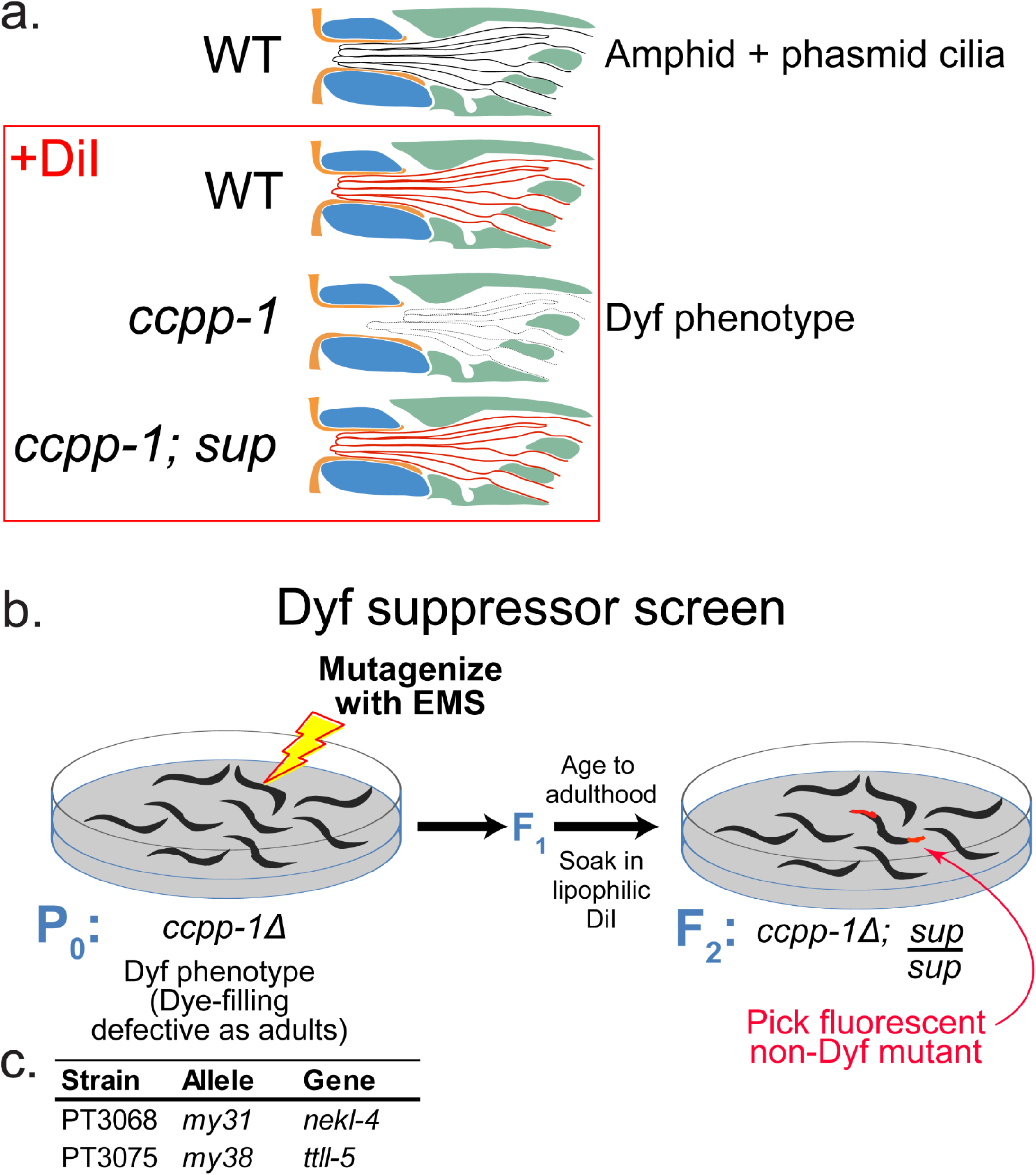
Suppressor screen design and *ccpp-1* Dyf suppressors identified. **a.** Illustration of the dye-filling assays. **b.** Diagram of EMS mutagenesis screen for *ccpp-1* Dyf suppressors (*sup*). **c.** Outcrossed single-gene suppressors identified from mutagenesis screen and sequenced. Names of strains and alleles are indicated, as well as the genes identified through sequencing.

We first tested candidate genes for suppression of the *ccpp-1* Dyf phenotype. Because *spas-1*/Spastin and *mei-1*/Katanin are implicated in the severing of glutamylated microtubules [27,28], we hypothesized that mutation in these genes might suppress *ccpp-1* hyperglutamylation-induced ciliary degeneration. Other candidates included the microtubule-depolymerizing kinesin-13 KLP-7 and the MAP kinase PMK-3, which interact with the deglutamylases CCPP-1 and CCPP-6 to regulate axon regrowth after laser-severing of *C. elegans* PLM touch receptor neurons [29]. Mutations in these and several other candidate genes failed to suppress the *ccpp-1* Dyf phenotype (**Supplemental Fig. 1**), suggesting that different molecules regulate microtubule stability in amphid and phasmid neuronal cilia.

To identify genes that act with *ccpp-1* to regulate ciliary stability, we conducted a F_2_ genetic screen for suppressors of the *ccpp-1(ok1821)* Dyf phenotype. We mutagenized >1000 P_0_ *ccpp-1* mutants, passaged to homozygose recessive mutations, performed dye-filling assays on F_2_ adults and screened for non-Dyf animals (**Fig. 1b**). Mutant lines were maintained and retested for suppression of the Dyf phenotype. From >100,000 mutagenized haploid genomes, we isolated 11 viable lines that displayed suppression of the *ccpp-1* Dyf phenotype when retested (**Supplemental Table 1**). The *my38* suppressor is a missense mutation in the *ttll-5* glutamylase gene (**Fig. 1c**) and the *my31* suppressor is a nonsense mutation in *nekl-4*, encoding a NIMA-related kinase (**Fig. 1c**; [30]).

### Single gene mutations in glutamylases *ttll-4, ttll-5,* and *ttll-11,* or the NIMA-related kinase *nekl-4*, suppress the *ccpp-1* dye-filling defect

Glutamylases of the tubulin tyrosine ligase-like TTLL family oppose CCP deglutamylase function [31]. The *C. elegans* genome encodes five TTLL glutamylases—*ttll-4, ttll-5, ttll-9, ttll-11,* and *ttll-15* [6,32]. *ttll-4* mutation suppresses the *ccpp-1* Dyf defect, suggesting that TTLL-4 glutamylates microtubules in the amphid and phasmid cilia [21]. We therefore tested deletion alleles in the other four *ttll* genes and found that deletion mutations in *ttll-5* and *ttll-11,* but not *ttll-9* or *ttll-15* (data not shown), suppressed the Dyf defect of *ccpp-1* (**Fig. 2a**).

**Figure 2.**
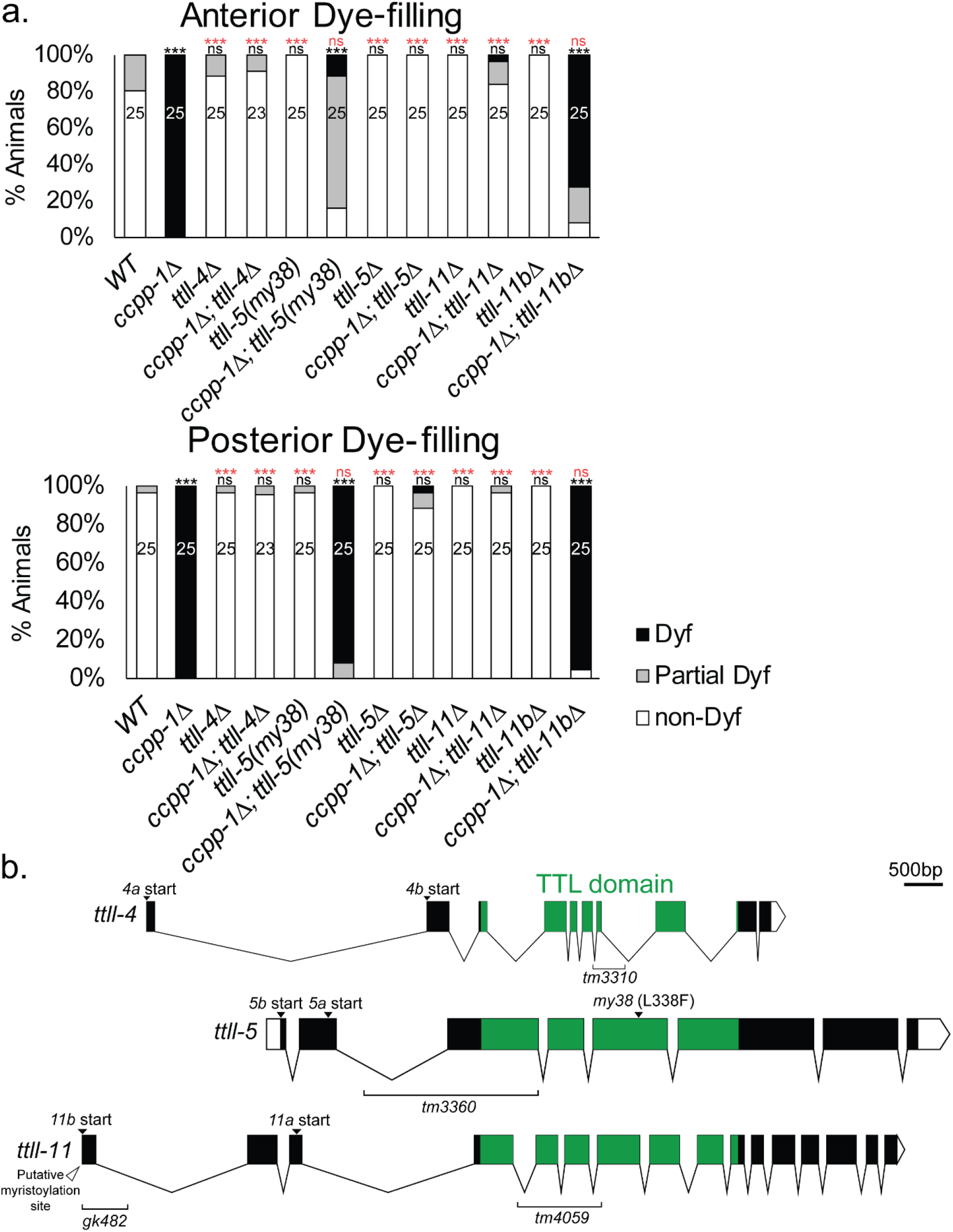
*ttll-4, ttll-5* and *ttll-11* act as suppressors of the *ccpp-1* Dyf phenotype. **a.** Suppression of *ccpp-1* Dyf phenotype by *ttll-4*, *ttll-5* and *ttll-11.* All strains marked with *Δ* are deletion alleles. *ttll-11bΔ* refers to the *ttll-11b(gk482)* allele. N of animals per strain is indicated on bars. *** indicates p ≤ 0.0001 by Kruskall-Wallis one-way ANOVA analysis and post hoc Dunn’s multiple comparison test. Black indicates significance relative to wild type and red indicates significance relative to *ccpp-1Δ*. **b.** Gene diagrams of *ttll-4, ttll-5* and *ttll-11,* aligned by TTL domain, including alleles used and important features.

The *ttll-4* gene encodes two protein isoforms by alternative splicing, TTLL-4A (601 AA) and TTLL-4B (563 AA) (**Fig. 2b;** [30]). Both isoforms contain a TTL domain (amino acid residues 138-476 in TTLL-4A; 100-438 in TTLL-4B) that is conserved among other TTLL genes. To study *ttll-4,* we used the *tm3310* allele, containing a deletion of 137 AAs of the TTL domain (**Supplemental Table 2**), which is predicted to reduce or eliminate glutamylase function. As previously shown [21], *ttll-4(tm3310)* strongly suppressed the *ccpp-1* Dyf phenotype (**Fig. 2a**).

The *ttll-5* gene also encodes two protein isoforms, TTLL-5A (677 AA) and TTLL-5B (730 AA) (**Fig. 2b, c;** [30]). The TTL domain in TTLL-5 spans AA residues 67-425 in TTLL-5a and 120-478 in TTLL-5b. The allele we identified in our screen, *my38,* encodes an L338F substitution, located in the TTL domain, that is predicted to affect ATP binding [A. Roll-Mecak, personal communication]. *ttll-5(my38)* suppressed *ccpp-1* in the amphids (**Fig. 2a**). We also characterized the recessive deletion allele *tm3360,* in which 49 AA are deleted, including 33 residues that align with the extended TTLL domain of mammalian TTLL5 [9,33]. Although this deletion is predicted to be in-frame, the large missing region of the TTLL domain might abolish or reduce substrate binding and glutamylase function [A. Roll-Mecak, personal communication]. *ttll-5(tm3360)* also strongly suppressed the *ccpp-1* Dyf phenotype in both amphids and phasmids (**Fig. 2a**).

TTLL-11 also has two isoforms, TTLL-11A (607 AA) and TTLL-11B (707 AA) (**Fig. 2b;** [30]). The TTL domain spans residues 24-388 in TTLL-11A and 124-488 in TTLL-11B. The *ttll-11(tm4059)* allele encodes an in-frame deletion in the TTL domain and is expected to disrupt function of both isoforms [22]. Like *ttll-4* and *ttll-5* mutations*, ttll-11(tm4059)* suppressed the *ccpp-1* Dyf phenotype in amphids and phasmids (**Fig. 2a**). In contrast, *ttll-11b(gk482)*, which deletes coding sequence from only the long B isoform, did not suppress *ccpp-1.* We cannot completely rule out the possibility that *gk482* reduces expression of both TTLL-11A and TTLL-11B. However, transcriptional GFP reporters suggest that TTLL-11B is expressed in extracellular vesicle-releasing neurons (EVNs) and not amphids [34], whereas TTLL-11A may be expressed in amphids [22]. These data suggest that TTLL-11A opposes the activity of CCPP-1 in amphid and phasmid neurons.

We isolated a *ccpp-1* Dyf suppressor, *my31,* which is a nonsense mutation in exon 5 of the *nekl-4* gene that produces NEKL-4(W199X) (**Supplemental Table 2**). *nekl-4(my31)* moderately suppressed *ccpp-1* Dyf in both amphid and phasmid neurons, and was non-Dyf as a single mutant (**Fig. 3a**). *nekl-4* encodes a 981 AA putative serine-threonine kinase (**Fig. 3b**). Prosite and InterPro scans detected a kinase domain (residues 453-725), and two armadillo (ARM) repeats (residues 200-281); ARM repeats are a series of alpha helices which can be indicative of protein-protein interactions [35]. Using EMBOSS epestfind [36], we also identified a PEST domain, important for proteolytic degradation (residues 748-764) [37]. NEKL-4 is a homolog of human NEK10, a protein required for ciliogenesis and associated with ciliopathies ([25,26]; **Fig. 3c**). NEK10 also contains ARM repeats (residues 137-348 and 387-471), a kinase domain (residues 519-791), and a PEST domain (residues 896-921). However, the NEK10 kinase domain is flanked by coiled-coil domains, and contains a putative tyrosine kinase active site not found in NEKL-4. NEKL-4 identity ranges from 29.41% (compared with NEK10 isoform 3) to 48.54% (NEK10 isoforms 5, 6 and 7). The NEK10 and NEKL-4 kinase domain is 52.50% identical by BLAST [38] (**Supplemental Fig. 2**).

**Figure 3.**
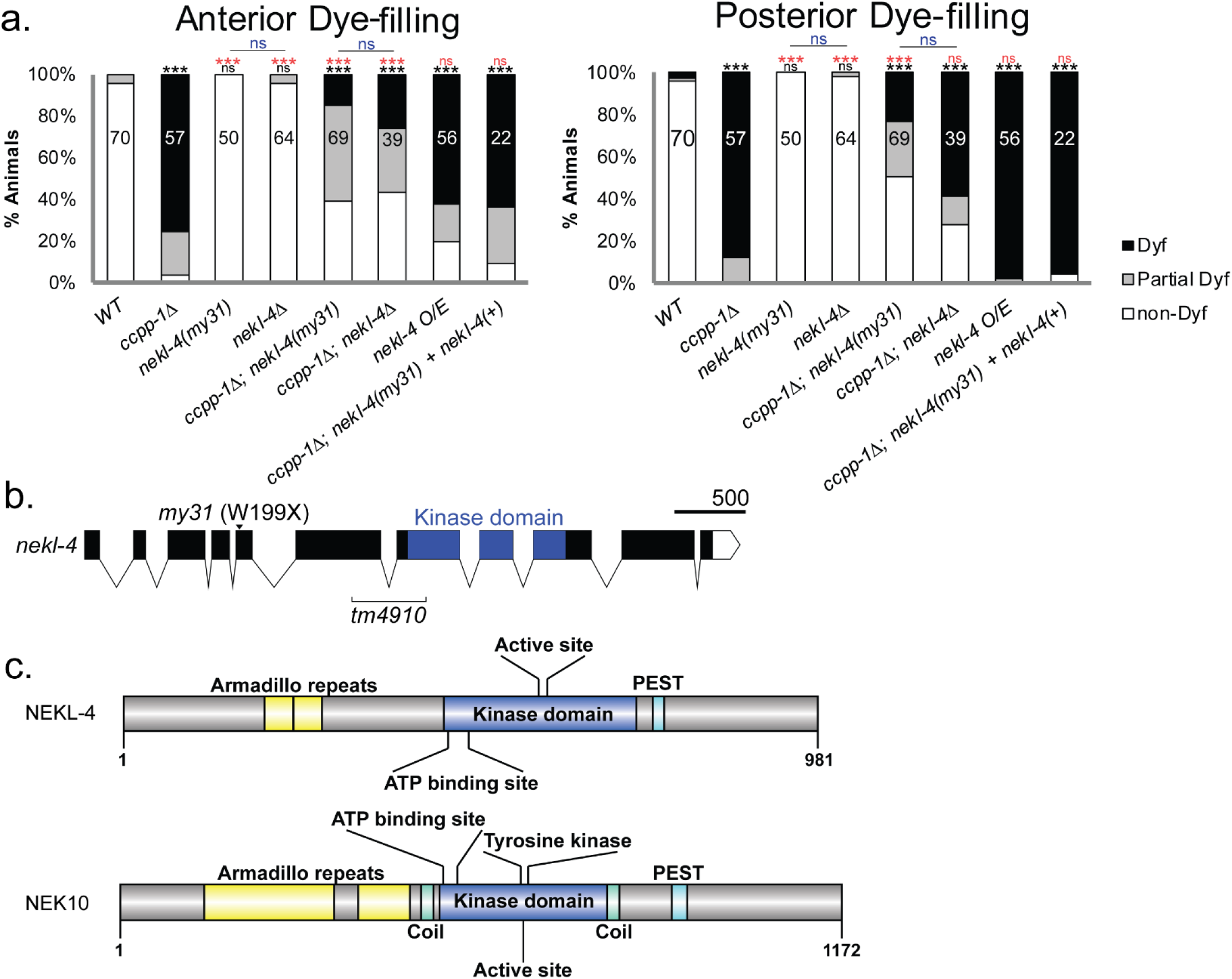
*nekl-4* suppresses the *ccpp-1* Dyf phenotype. **a.** Suppression of *ccpp-1* Dyf phenotype by *nekl-4*. N of animals per strain is indicated on bars. *** indicates p ≤ 0.0001 by Kruskall-Wallis one-way ANOVA analysis and post hoc Dunn’s multiple comparison test. Black indicates significance relative to wild type and red indicates significance relative to *ccpp-1Δ.* **b.** Diagram of *nekl-4* gene with alleles and important features indicated. **c.** Protein diagrams of NEKL-4 and its human homolog NEK10.

To confirm that *my31* was a mutation in the *nekl-4* gene, we performed transgenic rescue experiments by germline injection of the wild-type genomic copy of *nekl-4* (injected @1.5ng/μL) into *ccpp-1;nekl-4(my31)* mutants, and restored the *ccpp-1* Dyf phenotype (**Fig. 3a**). Injection of nearby genes *RO2F2.1* and *C27F2.1* failed to rescue the *ccpp-1* Dyf phenotype, further confirming *nekl-4(my31)* as the causative mutation (data not shown). We also characterized the deletion allele *tm4910,* which removes portions of exon 6 and exon 7 and replaces part of the kinase domain, including the putative ATP-binding lysine residue, with a glutamine residue. *nekl-4(tm4910)* single mutants are non-Dyf. *nekl-4(my31)* and *nekl-4(tm4910)* exhibited similar suppression of *ccpp-1* Dyf defects in amphid and phasmid neurons. Because both alleles were phenotypically indistinguishable, failed to complement for *ccpp-1* Dyf suppression (data not shown), and are likely loss-of-function alleles, we continued with the *nekl-4(tm4910)* deletion allele for ease of genotyping.

### NEKL-4 is expressed in the ciliated neurons but does not localize to cilia

We examined NEKL-4 expression pattern and subcellular localization using a transgenic extrachromosomal *nekl-4p∷nekl-4∷gfp* reporter and endogenously tagged *nekl-4∷mneongreen* and *nekl-4∷mscarlet* CRISPR reporters. Extrachromosomal NEKL-4∷GFP, driven by an endogenous promoter of 1609 base pairs before the start codon, localized in the axons, cell bodies, and dendrites of the ciliated neurons in the head of the worm, as well as the phasmid and PQR neurons in the tail, and was excluded from nuclei (**Fig. 4a-b**). NEKL-4∷GFP localized to the distal dendrite and the base of sensory cilia, but was not observed in the cilium. In the dendrites of some neurons, NEKL-4∷GFP accumulated in patches ranging from 2-5 μm in length. Overexpression of a *nekl-4p∷nekl-4∷gfp* transgene (injected at a concentration of 13 ng/μL) resulted in a Dyf phenotype (**Fig. 3d**), suggesting that levels or activity of NEKL-4 are important for function.

**Figure 4.**
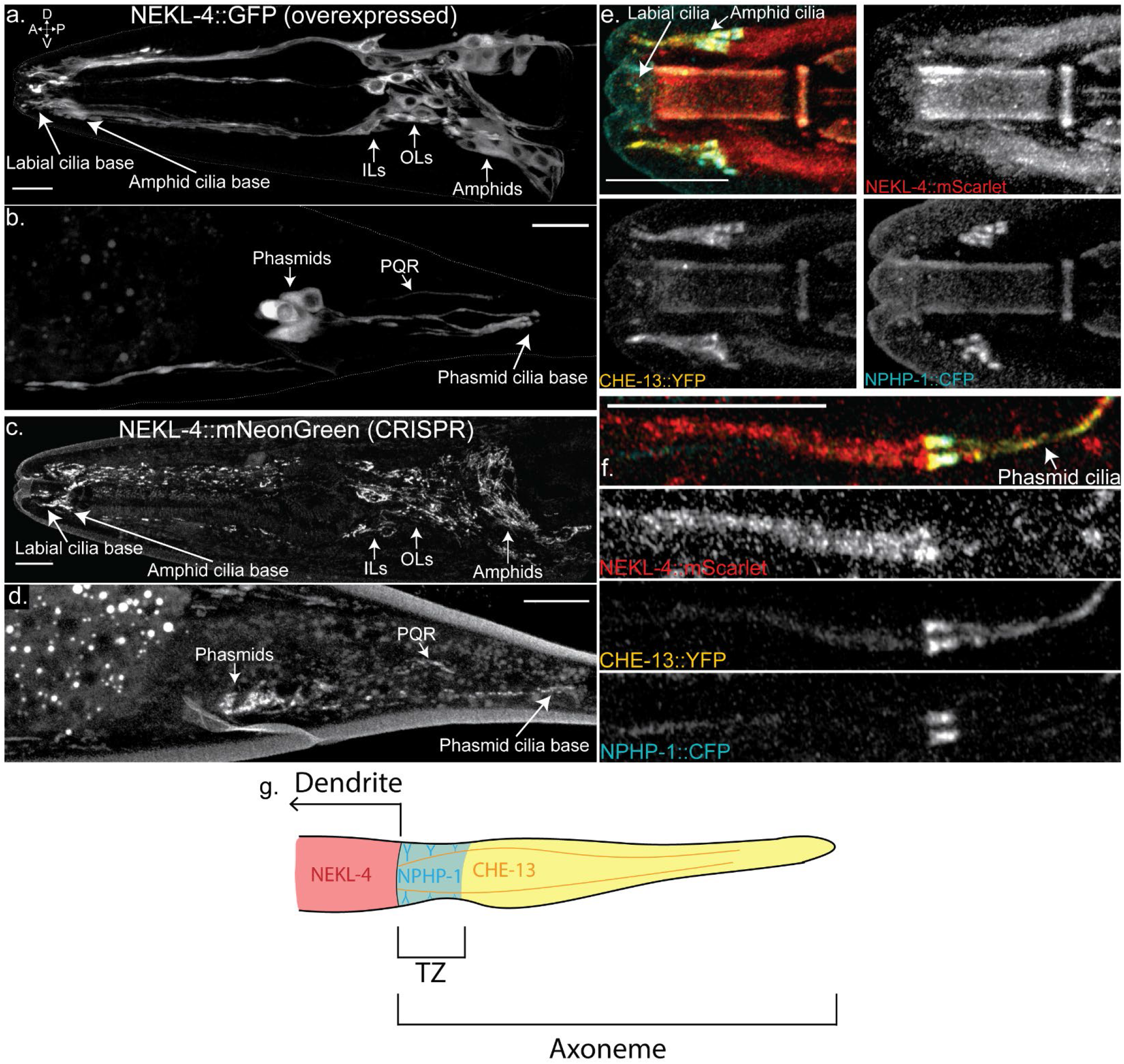
NEKL-4 is localized to ciliated neurons and is not enriched in the cilium. **a-b.** Localization of NEKL-4∷GFP, overexpressed by an extrachromosomal array. Scale = 10μm. **c-d.** Localization of NEKL-4∷mNeonGreen, expressed by inserting the fluorescent tag at the 3’ end of the endogenous *nekl-4* gene using CRISPR/Cas9. Scale = 10μm. **e-f.** Localization of NEKL-4∷mScarlet to the distal dendrite with NPHP-1∷CFP (transition zone) and CHE-13∷YFP (axoneme) as reference points. Scale = 10μm. **g.** Schematic of a phasmid cilium showing NEKL-4, NPHP-1, and CHE-13 localization.

We therefore examined the expression and localization of endogenous NEKL-4 by using CRISPR/Cas9 [39,40] to engineer endogenous *nekl-4∷mneongreen* and *nekl-4∷mscarlet* strains that did not exhibit a Dyf phenotype (data not shown) (**Fig. 4c-d**). These CRISPR-generated reporters use the *nekl-4* locus as well as a short flexible linker, and have significantly dimmer fluorescence than the extrachromosomal *nekl-4p∷nekl-4∷gfp* array. With both extrachromosomal and CRISPR reporters, NEKL-4 was observed in the same neurons and similar subcellular locations.

With the CRISPR-generated *nekl-4∷mneongreen* reporter, we were able to resolve fine structures in the cell bodies and dendrites. In the amphid cell bodies, NEKL-4∷mNG appeared as thin filaments. A similar pattern was seen in the outer labial (OL), inner labial (IL), and cephalic (CEP) cell bodies, though the filaments appeared to be thicker and brighter. The CRISPR-generated NEKL-4∷mNG reporter also localized in discrete, discontinuous patches in the dendrites. This pattern continued to the distal dendrite, terminating before entering the cilium of any neurons. We observed NEKL-4∷mNG movement in the distal dendrite (**Supplemental Movie 1**) and localization to the cell bodies and dendrites of the ray neurons in the male tail (**Supplemental Fig. 3**). The intricate localization pattern and movement of NEKL-4∷mNeonGreen suggests association with organelles or cytoskeletal components.

Because a *nekl-4∷gfp* reporter was previously reported [41] to localize in phasmid cilia, we examined colocalization of NEKL-4∷mScarlet with markers for the ciliary axoneme (*che-13∷yfp)* and transition zone (*nphp-1∷cfp*). NEKL-4∷mScarlet localization did not show significant overlap with NPHP-1∷CFP or CHE-13∷YFP, suggesting NEKL-4∷mScarlet localization was mainly restricted to the distal dendrite and periciliary membrane compartment (PCMC) (**Fig. 4e-f**). From these data, we conclude that NEKL-4 likely has a function in ciliated neurons, though the protein itself does not localize to the cilium (**Fig. 4g**).

### *ttll-4(tm3310)* and *ttll-5(tm3360)* suppress the *ccpp-1* glutamylation defect, and *ttll-11* activity is essential for glutamylation but not ciliogenesis

We examined glutamylation in the *ttll-4(tm3310), ttll-5(my38), ttll-5(tm3360), ttll-11(tm4059)*, *ttll-11b(gk482),* and *nekl-4(tm4910)* to determine their effects on microtubule glutamylation. We used the monoclonal antibody GT335, which recognizes the branch point of glutamate side-chains [42]. In wild type, GT335 labels the cilia of amphid, labial (OL, IL), and cephalic (CEP) neurons. Labial and cephalic cilia are seen at the anterior-most tip of the animal, surrounding the buccal cavity. Staining of labial and cephalic cilia appeared as singular axonemes [43]. In amphid channel cilia, GT335 specifically stains the doublet region middle segments that appear as two triangular bundles located posterior to the labial and cephalic cilia [43]. In wild type animals, GT335 strongly stained labial and cephalic cilia and amphid ciliary middle segments (**Fig. 5a**). In the *ttll-4(tm3310)* mutants, we observed a reduction in the percentage of animals with amphid, labial and cephalic staining, indicating TTLL-4 affects glutamylation (**Fig. 5d**). In the *ttll-5(my38)* and *ttll-5(tm3360)* single mutants, we observed normal GT335 staining. The *ttll-11(tm4059)* mutation, which affects both TTLL-11 isoforms, abolished GT335 staining, whereas *ttll-11b(gk482)* showed normal GT335 staining in amphid middle segments. We found that *nekl-4(tm4910)* displayed a normal GT335 staining phenotype, suggesting that NEKL-4 does not play a role in microtubule glutamylation. Our data suggests that TTLL-11 but not TTLL-4 or TTLL-5 is essential for initiating branch point glutamylation.

**Figure 5.**
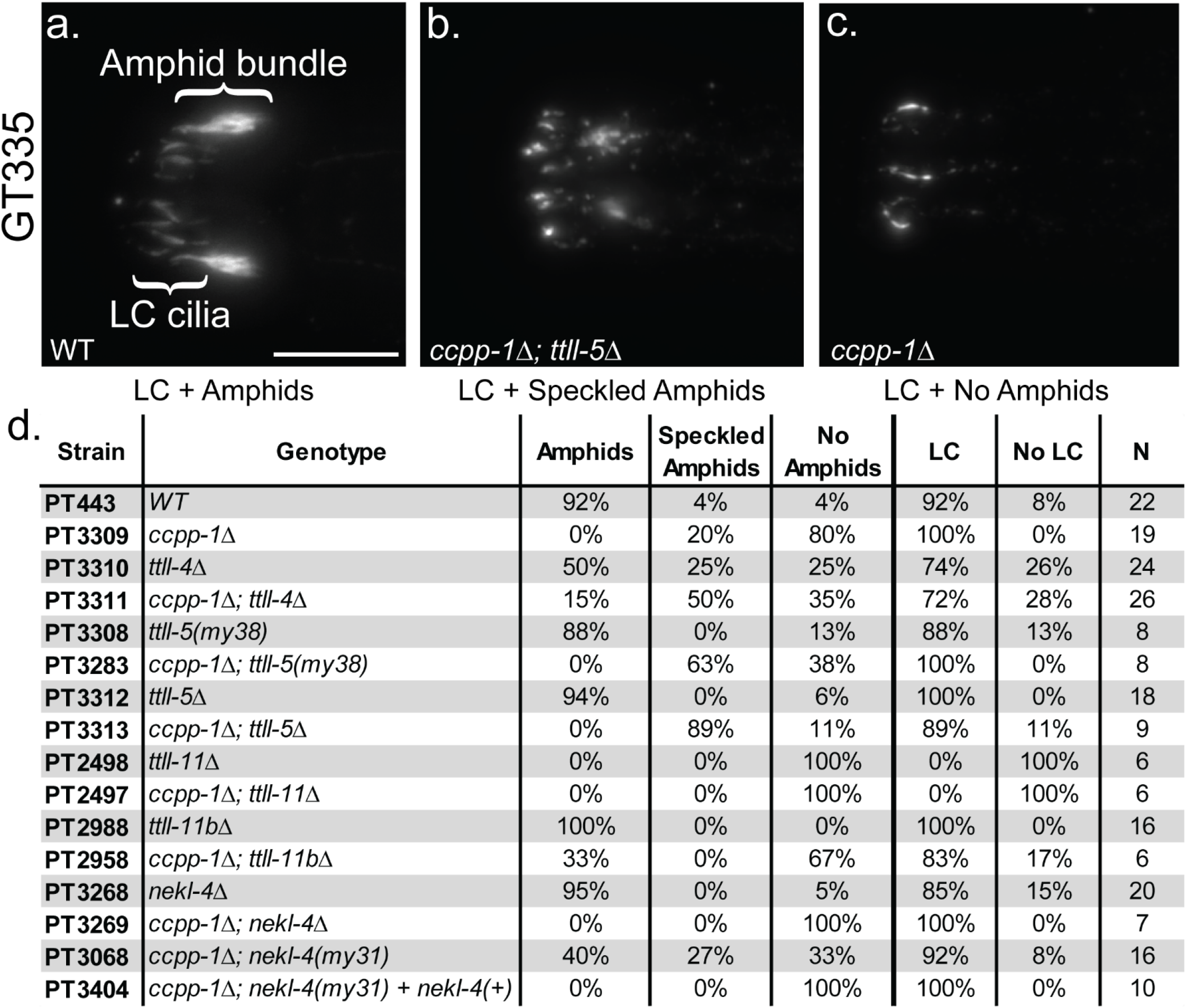
Effects of mutations on glutamylation of amphid, labial and cephalic (LC) cilia. **a-c.** Examples of GT335 antibody staining phenotypes. Scale = 10μm. **d.** Percentage of worms with GT335 stained amphids and/or labial and cephalic (LC) cilia. PT3404 was rescued with a *nekl-4p∷nekl-4∷gfp* plasmid @ 1.5ng/μL.

We next examined the effect of the suppressor mutations on GT335 staining in *ccpp-1* mutants. We analyzed both the presence of GT335 staining and the appearance of staining to examine subtle differences in glutamylation patterns. We observed a variation in the GT335 staining pattern of some mutants: the amphids were stained but the pattern was abnormal. We classified this staining pattern as a “speckled amphid” phenotype (**Fig. 5b**). We also observed abnormal phenotypes in ciliary glutamylation in *ttll* mutants and suppressor strains (**Fig. 5b-c**). In some mutants including *ccpp-1*, labial and cephalic cilia stained while amphid cilia did not, consistent with cell-specific regulation of the Tubulin Code [2,22,29,44] (**Fig. 5c**).

In *ccpp-1* mutants, amphid channel cilia GT335 staining was often absent or occasionally speckled, may reflect nearly complete ciliary microtubule degradation [21] (**Fig. 5d**). Speckled staining might indicate that ciliary microtubules are only partially degraded. In *ttll-4(tm3310),* 50% of animals had speckled or absent amphids. We observed a restoration of GT335 staining in amphid channel cilia in 15% of the *ccpp-1; ttll-4* double mutant, consistent with suppression of ciliary degeneration (**Fig. 5d**). Although *ttll-5(my38)* and *ttll-5(tm3360)* single mutations had no effect on GT335 staining of amphid channel cilia, *ccpp-1; ttll-5* double mutant amphid cilia had a higher incidence of speckled middle segments compared to the *ccpp-1* single mutant.

*ttll-11(tm4059)* single and *ccpp-1; ttll-11(tm4059)* double mutants showed no GT335 staining, yet both strains were non-Dyf (**Fig. 5d, Fig. 2a**). *ccpp-1;ttll-11b(gk482)* mutants, in contrast, had partial restoration of amphid and labial and cephalic GT335 staining. We suggest that *ttll-11* activity is required for glutamylation to occur, and therefore, we cannot conclude that the absence of GT335 staining indicates an absence of microtubules, only an absence of glutamylation. Combined, our data indicates that reducing glutamylation suppresses microtubule degeneration due to loss of CCPP-1 function and that glutamylation is not essential for ciliogenesis.

Unlike *ttll-4* and *ttll-5* mutations*, nekl-4(tm4910)* did not suppress the abnormal GT335 staining in *ccpp-1.* Curiously, although *nekl-4(my31)* and *nekl-4(tm4910)* phenocopied each other in *ccpp-1* Dyf suppression, we observed differences in their effects on *ccpp-1* GT335 phenotype. *nekl-4(my31)* but not *nekl-4(tm4910)* showed partial suppression of the *ccpp-1* GT335 phenotype (**Fig. 5d**). We hypothesize that *nekl-4(my31)* could be a stronger suppressor than *nekl-4(tm4910)* because it encodes an early stop codon that eliminates the entire kinase domain. In transgenic animals, introduction of a *nekl-4(+)* genomic construct rescued *ccpp-1;nekl-4(my31)* amphid GT335 staining to appear similar to *ccpp-1* alone (Fig. 5d). We conclude that NEKL-4 may act downstream or independent of CCPP-1-mediated deglutamylation.

### *ttll-5* and *ttll-11* suppress heterotrimeric kinesin-2 span shortening in *ccpp-1* mutants

*C. elegans* amphid and phasmid cilia are built by the cooperative action of heterotrimeric kinesin-2 and homodimeric OSM-3/KIF17 [45,46]. Heterotrimeric kinesin-2 travels along the doublet-containing middle segment region, which makes up approximately half of the length of the amphid channel and phasmid cilia and correlates with GT335 staining [45,46]. We therefore examined the middle segment localization of KAP-1∷GFP, the non-motor subunit of heterotrimeric kinesin-2 [45]. In wild-type phasmid cilia, the span of KAP-1∷GFP was approximately 4 μm (**Fig. 6a**). In *ccpp-1* mutants, the KAP-1∷GFP span was significantly reduced to ~2 μm. In *osm-3(p802)* mutants that lack distal singlet microtubules, we observed a similar shortened KAP-1∷GFP span of 2 μm [18,20,43]. Consistent with the non-Dyf phenotype of *ttll-4, ttll-5, ttll-11,* or *nekl-4(tm4910)* single mutants, KAP-1∷GFP spans were similar to wild type (**Fig. 6a**). Mutations in *ttll-5* and *ttll-11* that suppressed the Dyf phenotype of *ccpp-1* also suppressed the KAP-1∷GFP span shortening of *ccpp-1* (**Fig. 6a**). *nekl-4(tm4910)* did not suppress the *ccpp-1* KAP-1∷GFP span shortening (**Fig. 6a**). We considered the following possibilities to explain the shortened KAP-1∷GFP span in *ccpp-1*: (i) microtubule loss in *ccpp-1* mutants, or (ii) altered association with OSM-3 in the ‘handover zone’, which may be needed for heterotrimeric kinesin-2 to travel the appropriate distance [47]. Both these possibilities are consistent with roles for the Tubulin Code in regulating microtubule stability and motor properties. Our data on the *ttll* gene family of suppressors indicate a role for changes in microtubule glutamylation levels to contribute to rescue of CCPP-1 loss of function; this rescue could occur either by suppressing ciliary microtubule loss, altering motor protein-IFT particle interactions, or both. Additionally, our data on *nekl-4(tm4910)* reveals the presence of a mechanism for *ccpp-1* Dyf suppression that does not involve changes in microtubule glutamylation levels or heterotrimeric kinesin-2 location.

**Figure 6.**
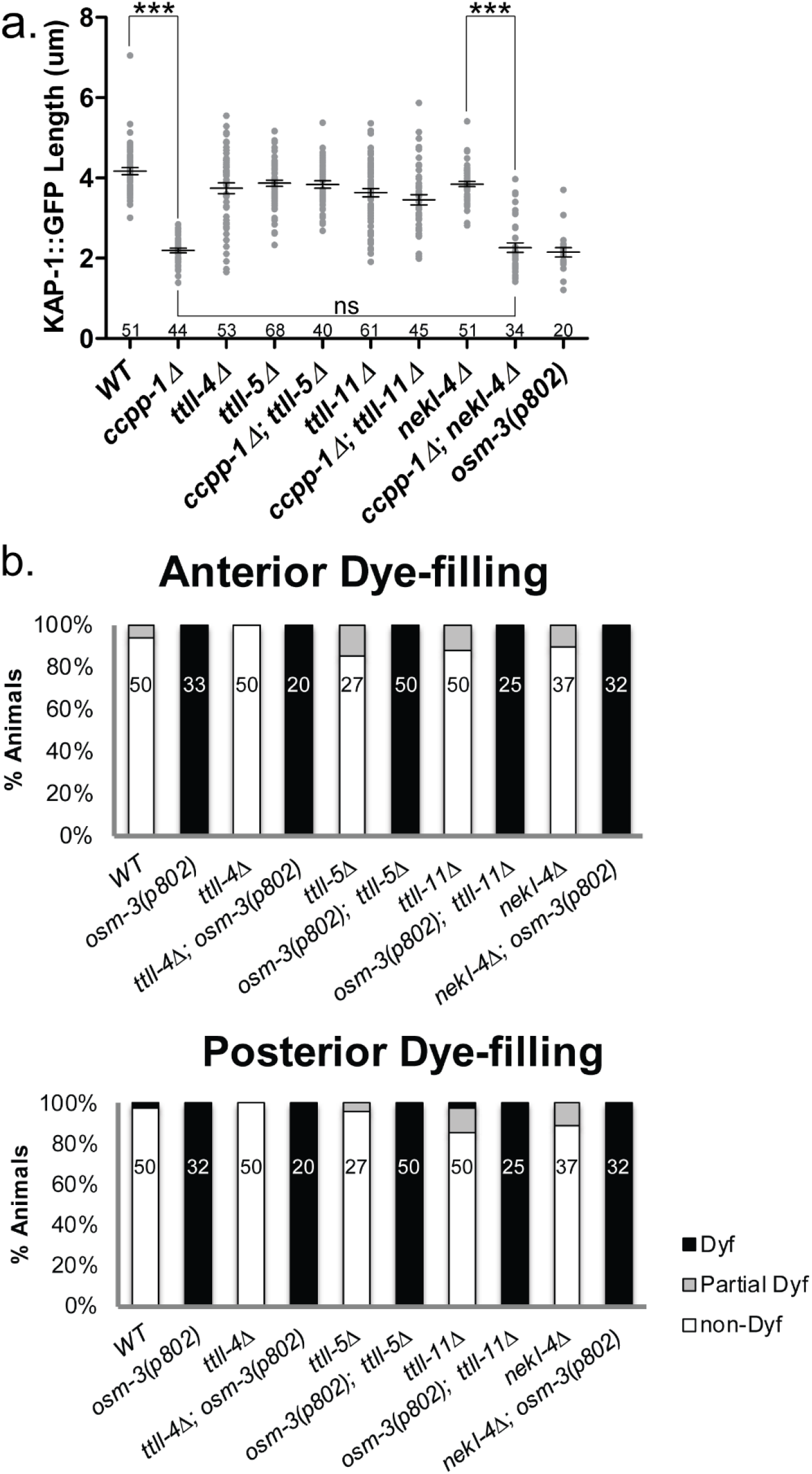
Mutations in TTLL genes suppress *ccpp-1* defects in KAP-1 middle segment localization, but do not suppress distal segment defects in *osm-3(p802)* mutants. **a.** Length of KAP-1∷GFP span in the phasmid neurons. Mean ± SEM. *** indicates p ≤ 0.0001 by Kruskall-Wallis one-way ANOVA analysis and post hoc Dunn’s multiple comparison test. n of cilia per strain is indicated. **b.** Dye-filling assay showing lack of *osm-3(p802)* suppression by *ttll-4, ttll-5, ttll-11* or *nekl-4.* N of animals for each strain is indicated.

In addition to heterotrimeric kinesin-2, homodimeric OSM-3/KIF17 kinesin-2 is important for ciliogenesis in amphid and phasmid neurons [18,48,49]. Previous studies indicated that *nekl-4* genetically interacts with *osm-3* [41]. *nekl-4(cas396)*, which affects the kinase domain, partially suppresses the Dyf phenotype and distal segment loss of *osm-3(sa125)* mutants [41]. The *osm-3(sa125)* G444E missense mutation is proposed to relieve the autoinhibition of the coiled-coil stalk of OSM-3 by fixing its hinge region in an extended conformation, but severely impairs its motility. OSM-3(G444E) can still associate with heterotrimeric kinesin-2 [41,49]. We examined genetic interactions between the null alleles *nekl-4(tm4910)* and *osm-3(p802)* allele, which phenocopies the *osm-3(sa125)* amphid cilia truncation phenotype [18,41]. The *osm-3(p802)* null mutation was not suppressed by *nekl-4(tm4910)*, indicating that interactions between *nekl-4* and *osm-3* are allele-specific, and may depend on the physical or functional interactions between OSM-3 and NEKL-4 (**Fig. 6b**).

## Discussion

We used a forward genetic screen and candidate approach to identify genes that act with the *ccpp-1* deglutamylase to control ciliary microtubule stability. We identified tubulin code writers and effectors of ciliary stability as modulators of degenerative phenotypes associated with CCPP-1 loss. Our data provides insight into how glutamylation is precisely controlled to maintain ciliary structure and function and to prevent neurodegeneration. *ccpp-1* mutants can be rescued by reducing microtubule glutamylation levels through the loss of function of tubulin glutamylating enzymes TTLL-4, TTLL-5, or TTLL-11. Our findings are in agreement with studies in *Chlamydomonas* where reducing microtubule glutamylation has a stabilizing effect on axonemal defective mutants ([50,51]; for a summary of phenotypes refer to **Supplementary Table 3**).

The NIMA related kinase NEKL-4 opposes CCPP-1 action and may act independent of microtubule glutamylation state. Loss of *nekl-4* function suppresses the *ccpp-1* Dyf phenotype while animals overexpressing NEKL-4∷GFP are Dyf. Our findings are consistent with previous literature on the roles of NIMA kinases in regulating axonemal stability [51–53]. Since *nekl-4(tm4910)* does not affect GT335 staining, it is unlikely that NEKL-4 regulates a glutamylase or deglutamylase. NEKL-4 is also unlikely to be a reader of ciliary glutamylation because it is largely absent from cilia. We speculate that other molecules might function in a pathway that reads out the ciliary MT glutamylation information and relay it to NEKL-4, which then activates proteins that directly or indirectly lead to MT severing or depolymerization by phosphorylating them. Alternatively, NEKL-4 and the glutamulation machinery may act in parallel pathways to regulate ciliary stability (**Fig. 7**).

**Figure 7.**
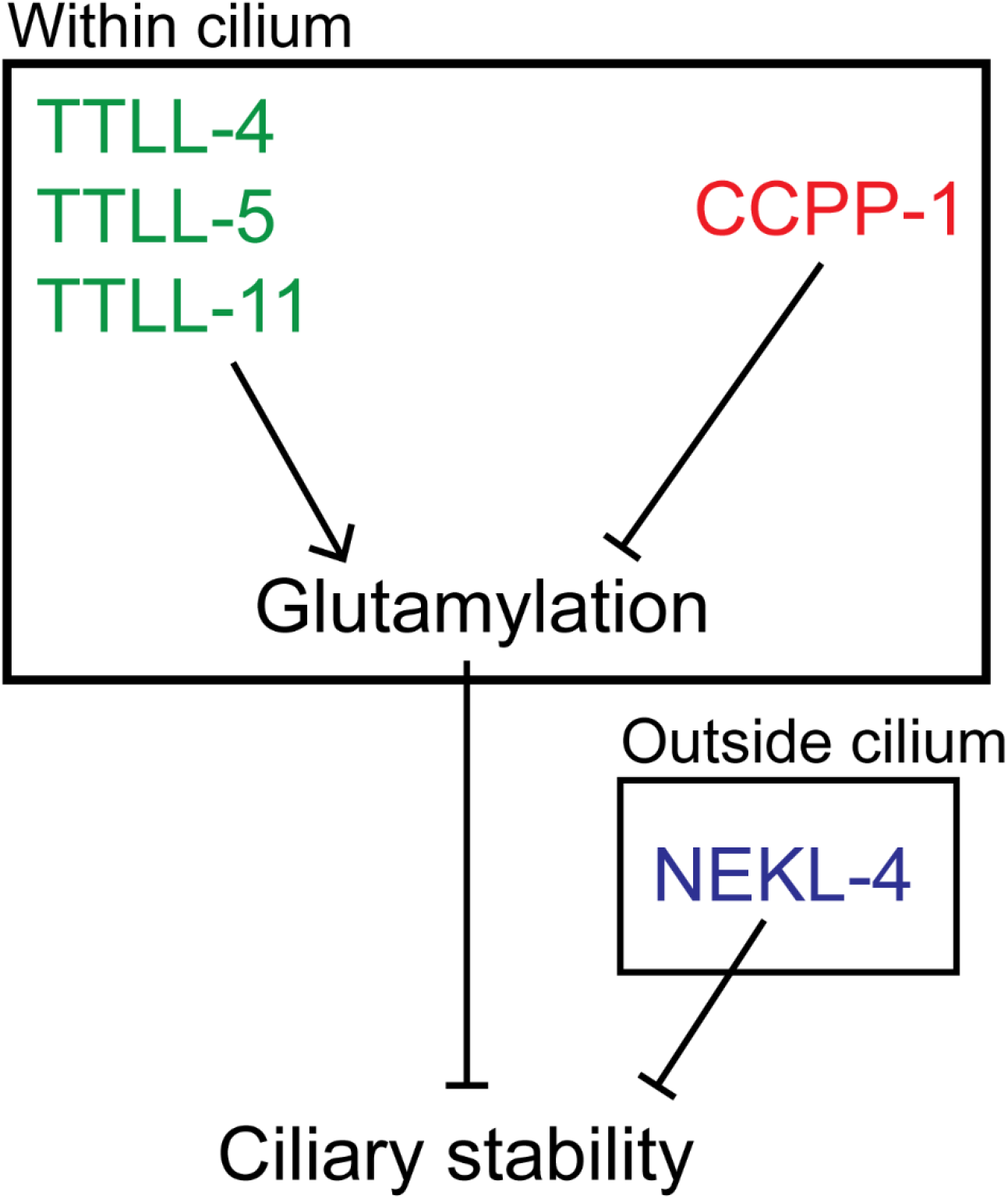
Schematic of TTLL-4, TTLL-5, TTLL-11, CCPP-1, and NEKL-4 interactions and effects on ciliary stability.

What could be the mechanism of rescue of *ccpp-1* loss of ciliary stability? Our data here eliminated a role for microtubule severing by katanin (MEI-1) and spastin (SPAS-1) in modulating the loss of ciliary stability in *ccpp-1* mutants. Our results also did not support a role for other known effectors of microtubule glutamylation— MAP kinase PMK-3 and microtubule-depolymerizing kinesin-13 (KLP-7), in the degradation of ciliary microtubules in the absence of CCPP-1 function. Therefore, other unknown molecules and pathways likely sense hyperglutamylation and mediate the destruction of ciliary microtubules in amphid and phasmid neurons. We put forth the idea that rescue of *ccpp-1* loss of ciliary stability might involve changes in microtubule dynamics. Hypo-glutamylation due to the loss of TTLL-4, TTLL-5, or TTLL-11 function may reduce tubulin turnover in *ccpp-1* mutant neurons and stabilize axonemal microtubules (refer to model in **Fig. 7**). This hypothesis is supported by previous studies in *Chlamydomonas* that revealed a link between the levels of tubulin polyglutamylation and the rates of tubulin turnover [54]. Conversely, *nekl-4(tm4910)* mediated rescue of *ccpp-1* Dyf might involve an *increase* in tubulin turnover. This assertion is supported by studies on a NIMA-related kinase *CNK11* in *Chlamydomonas reinhardtii. cnk11* mutants, similar to *C.elegans nekl-4* mutants, rescue mutants with ciliary instability. *cnk11* mutants have an increase in tubulin turnover that depends on the glutamylation state of ciliary microtubules [51]. Although it is not entirely clear how an *increase* in tubulin turnover rate could rescue the degradative effects of hyperglutamylation, the progressive nature of the Dyf phenotype of *ccpp-1* mutants indicates that the degeneration of hyperglutamylated microtubules is a gradual process. Whether this lag could be attributed to the time it takes for the amphid/phasmid ciliary microtubules to excessively elongate glutamate side chains that triggers degeneration is unknown.

Recent studies have ascribed a role for NEKs in regulating endocytosis. *C. elegans* NEKL-2 and NEKL-3 control clathrin-mediated endocytosis and trafficking in non-ciliated cells [55–57]. NEKL-4 localization to the periciliary membrane compartment - a site of endocytosis that regulates IFT, ciliary membrane volumes, and ciliary protein levels - hints at a role in endocytic trafficking at the ciliary base [58]. Reduction in endocytosis can rescue the Dyf defect of a *gpa-3*QL mutant [59]. Our data leaves open a role for NEKL-4 in endocytosis, and the possibility that *nekl-4* rescues *ccpp-1* ciliary degeneration through modulation of endocytic activity. The mechanisms of NEKL-4 action remain unknown and are exciting future directions.

NEK10 potential substrates include kinesins, IFT components, and other ciliary proteins that have homologs in *C. elegans* ([26,60]**; Supplementary Table 4**). In human bronchial epithelial cell (HBEC) culture, NEK10 mutation caused a reduction in phosphopeptides of gene products including KIF13B, KIF19, IFT140, IFT46, IFT88, and MAK [26]. These genes and their *C. elegans* orthologs *klp-4, klp-13, che-11, dyf-6, osm-5*, and *dyf-5* are involved with microtubule dynamics, IFT, and ciliary structure [41,61–69]. Therefore, NEKL-4 may act on IFT proteins or other ciliary proteins.

Ciliary microtubules are heavily glutamylated and a growing list of reports links hyper- or hypo-glutamylation to disease. The genes identified herein are associated with human disorders. Loss of CCP1 causes retinal dystrophy, and the infantile-onset neurodegeneration of cerebellar neurons, spinal motor neurons, and peripheral nerves [70,71]. Glutamylation may also play an important role in ciliopathies including Joubert Syndrome [72] and hereditary retinal degeneration [24]. NEK kinases are also implicated in ciliopathies [26,73] and cancer [74–76]. Our work suggests that NEK kinases may represent new targets for treatment of human neurodegenerative and ciliopathic disease, and that insights into the function of NEKs might be crucial for understanding pathways by which the Tubulin Code specializes microtubule networks and ciliary integrity in human ciliopathies and neurodegenerative disorders.

## Materials and Methods

### *C. elegans* strains and maintenance

Nematodes were cultured on Nematode Growth Media (NGM) agar plates containing a lawn of OP50 *E. coli* and incubated at 20°C, as previously described [77]. Strains used are listed in **Supplemental Table 4**. Some strains contained *him-5(e1490)* and/or *myIs1[pkd-2∷gfp+unc-122p∷gfp] pkd-2(sy606),* which were considered wild type.

### *ccpp-1* EMS mutagenesis screen

To identify the pathway by which *ccpp-1* mutations cause the Dyf phenotype [21], we performed a forward genetic screen for suppressors of the Dyf defect of *ccpp-1* mutants. Ethyl methanesulfonate (EMS; Sigma Aldrich Cat. M0880) was used to mutagenize *ccpp-1(ok1821)* mutants {either Strain PT2168--*ccpp-1(ok1821); myIs1[pkd-2∷gfp + unc-122p∷gfp] pkd-2(sy606); him-5(e1490)* or OE4160--*ccpp-1(ok1821)*}. L4 larvae were suspended in 1 ml of M9 buffer [78] with a final concentration of 47 mM EMS for four hours, before washing three times in M9 and depositing worm suspension onto an NGM plate seeded with OP50 *E. coli* to recover. We then transferred 10 or 20 of these mutagenized P_0_ hermaphrodites to new NGM plates and allowed them to lay eggs on 100mm NGM plates seeded with OP50 for one day or two days before removing them. When the F_1_ hermaphrodites reached adulthood, we synchronized the age of F_2_ progeny by hypochlorite treatment of gravid adult F_1_ animals [78] and replating embryos on NGM plates with OP50. We then incubated plates for four days at 20°C or five days at 15°C (to allow the age-synchronized F_2_ progeny to reach adulthood) before washing them off plates using M9 to be dye-filled using the method described below. *ccpp-1* mutants without suppressor mutations typically are completely Dye-filling defective (Dyf) by adulthood. Therefore, to isolate suppressor mutants, we picked F_2_ adult animals that were non-Dyf or incompletely Dyf. We isolated up to five suppressor mutant isolates corresponding to a single P_0_ plate to allow for some lethality, but in most cases kept only a single suppressor from each P_0_ plate, except as noted in **Supplemental Table 1**.

In total, we examined more than 100,000 haploid genomes. We isolated 93 non-Dyf or partially Dyf F_2_ animals, of which 79 either died, were sterile, or were Dyf (not suppressed) in a subsequent secondary dye-filling assay to confirm the phenotype. 14 isolates were viable and produced non-Dyf or partial Dyf progeny in secondary screens, of which we consider 11 to be independent isolates (**Supplemental Table 1**).

### Whole-genome sequencing

To facilitate identification of suppressor mutations, whole-genome sequencing (WGS) was performed using the strategy described in Doitsidou *et al* [79]. Mutants were crossed with PT3037, a strain in which *ccpp-1* had been introgressed into the Hawaiian wild isolate genomic background by backcrossing ten times into CB4856. From crosses of PT3068 (*my31*) X PT3037, at least 20 suppressed F_2_s were isolated; these F_2_s were pooled for WGS to identify genomic intervals that lacked PT3037-derived single-nucleotide polymorphisms (SNPs); novel SNPs within those intervals potentially represent the suppressor mutation.

Adapter-ligated libraries were prepared from genomic DNA using the manufacturer’s protocol (Illumina, San Diego, CA) from PT3037 and the pooled suppressed F_2_s. A HiSeq 2000 instrument (Illumina, San Diego, CA) was used to sequence the genomic libraries. To identify the *my31* mutation, we used a pipeline described in Wang *et al* [80]. Briefly, the pipeline was as follows: BFAST for alignment [81], SAMtools for variant calling [82], and ANNOVAR for annotation [83]. We compared sequence data with *C. elegans* genome version WS220 to identify mutations. To identify mapping intervals that might contain the *my31* suppressor, we plotted Hawaiian SNP density against chromosome position. Non-Hawaiian SNPs that were also identified in PT3037 were removed as background. Candidate mutations for *my31* were identified as novel homozygous non-synonymous coding variants within the mapping interval.

### Protein domain identification, homology analysis and alignments

Sequences of *ttll-4, ttll-5, ttll-11,* and *nekl-4* were obtained from WormBase [30] and protein sequences were obtained from WormBase or UniProt [85]. Structural predictions and protein domain information were generated with InterPro [86] and ScanProsite [87]. The PEST proteolytic degradation site in NEKL-4 was identified using EMBOSS epestfind [36]. NEKL-4 homology to NEK10 was predicted using protein BLAST [88] and human NEK10 and *C. elegans* NEKL-4 kinase domains were aligned using T-COFFEE Expresso [33]. All scans and queries described above were performed using default settings. Gene diagrams were created with the WormWeb Exon-Intron graphic maker [89], and protein diagrams were created with DOG 2.0 software [90,91]. The NEK10/NEKL-4 kinase domain alignment figure was generated with BoxShade [92] from the T-COFFEE sequence alignment.

### Dye-filling assays

Three days prior to performing dye-filling assays, healthy, non-crowded plates containing many gravid hermaphrodite animals were washed and bleached to synchronize the worms. Day 1 and Day 2 adults were washed from the plate with 1mL M9 buffer and transferred to 1.5mL tubes. A stock solution of 2.5mg/mL DiI (Thermo Fisher Cat. D282) was added to a 1:1000 dilution and worms were incubated with gentle shaking for 30 min at room temperature. Prior to scoring, worms were briefly spun at 2000RPM and plated on new seeded NGM plates for approximately 1hr to excrete excess dye. The presence and intensity of dye in the amphid and phasmid neurons was visually scored using a dissecting microscope. Worms scored as non-Dyf were similar in brightness to wild type, and Dyf worms had no staining. Partial Dyf worms had dye present, but distinguishably less bright than wild type. Kruskall-Wallis one-way ANOVA analysis (where non-Dyf=2, partial Dyf=1. and Dyf=0) and posthoc Dunn’s multiple comparison test were performed in Prism (Graphpad Software).

### Antibody staining

We performed immunofluorescence microscopy using the protocol described in O’Hagan *et al* [22]. We synchronized animals by bleaching and fixed as Day 1 adults. Fixation was accomplished by washing animals from 3 NGM plates using M9 buffer, then washing animals in a 15 ml conical tube 3 more times with M9 over one hour. Worms were chilled on ice before washing in ice-cold Ruvkun buffer (80 mM KCl, 20 mM NaCl, 10 mM EGTA, 5 mM spermidine-HCl, 15 mM Pipes, pH 7.4 and 25% methanol) plus 2% formaldehyde in 1.5ml centrifuge tubes. The tubes were immersed in liquid nitrogen and melted under tap water to crack the worms’ cuticles. Worms were then washed twice with Tris-Triton buffer (100 mM Tris-HCl, pH 7.4, 1% Triton X-100 and 1 mM EDTA), suspended in Tris-Triton buffer+1% β-mercaptoethanol, and incubated overnight at 37°C. The next day, worms were washed with 1X BO_3_ buffer (50 mM H_3_BO_3_, 25 mM NaOH) + 0.01% Triton, and suspended in 1X BO_3_ + 0.01% Triton buffer + 10 mM DTT for 15 minutes with gentle agitation at room temperature. Worms were then washed with 1X BO_3_ buffer (50 mM H_3_BO_3_, 25 mM NaOH) + 0.01% Triton, and suspended in 1X BO_3_ + 0.01% Triton buffer + 0.3% H_2_O_2_ for 15 minutes with gentle agitation at room temperature. After washing once with 1X BO_3_ + 0.01% Triton buffer, worms were washed for 15 minutes in Antibody buffer B (1X PBS, 0.1% BSA, 0.5% Triton X-100, 0.05% sodium azide, 1mM EDTA) with gentle agitation at room temperature. Fixed worms were stored in Antibody buffer A (1X PBS, 1% BSA, 0.5% Triton X-100, 0.05% sodium azide, 1mM EDTA) at 4°C for up to one month before antibody staining. Animals were stained overnight at room temperature with a 1:450 dilution (in Antibody Buffer A) of GT335 (Adipogen Cat. AG-20B-0020-C100), a monoclonal antibody which binds the branch point of both monoglutamylated and polyglutamylated substrates [42]. Stained worms were washed with several changes of Antibody B Buffer with gentle agitation at room temperature over several hours. After rinsing with Antibody Buffer A, Alexa-fluor 568-conjugated donkey anti-mouse secondary antibody (Invitrogen Cat. A10037) was added at a final dilution of 1:2500 and incubated for 2 hours at room temperature with gentle agitation. Worms were then washed with several changes of Antibody Buffer B over several hours before mounting on 2% agarose pads for imaging.

We observed variation between and within individual GT335 staining experiments with respect to pattern and intensity. Environmental factors may cause changes in levels of ciliary glutamylation as measured by GT335 intensity [93]. It is therefore possible that small variations in experimental techniques or day-to-day lab conditions could have impacted our results. However, the GT335 staining phenotype of each mutant was generally consistent across trials.

### Confocal imaging

Day 1 adult hermaphrodites were anaesthetized with 10 mM levamisole and mounted on 4% agarose pads for imaging at room temperature. Confocal imaging was performed with a Zeiss LSM 880 inverted microscope with an Airyscan superresolution module using a LSM T-PMT detector and ZenBlack software (Carl Zeiss Microscopy, Oberkochen, Germany). Laser intensity was adjusted to avoid saturated pixels. Images were acquired using a 63x/1.4 Oil Plan-Apochromat objective in Airyscan Fast mode and deconvolved using Airyscan processing. Image files were imported into Fiji/ImageJ [94] with the BioFormats Importer plugin for linear adjustment of contrast and creation of maximum intensity projections, and Adobe Photoshop CS5 was used to trace the outlines of the worm if needed. Images were placed in Adobe Illustrator CS5 for figure assembly.

### KAP-1∷GFP localization analysis

Nematodes were mounted as above. Widefield images were acquired on a Zeiss Axio Observer with Colibri 7 LEDs and ZenBlue software (Carl Zeiss Microscopy, Oberkochen, Germany) using a Photometrics Prime 95B sCMOS camera (Teledyne Photometrics, Tucson, AZ). A 100x/1.4 Oil Plan-Apochromat objective was used for imaging the phasmid cilia of Day 1 hermaphrodite animals. Images were imported into Fiji/ImageJ [94] and the length of KAP-1∷GFP labeling was measured for each cilium using maximum intensity projections. Length was measured starting at the transition zone, which is distinct and bright, and ending when fluorescence was no longer visible. Kruskall-Wallis one-way ANOVA analysis and posthoc Dunn’s multiple comparison test were performed in Prism (Graphpad Software).

We also wanted to examine if *ttll-4* suppressed the *ccpp-1* KAP-1∷GFP span defect, but were unable to obtain a viable *ccpp-1;ttll-4; kap-1∷gfp* strain, possibly due to synthetic lethality with the overexpressed *kap-1∷gfp* reporter.

### Plasmid construction

For creation of pRO139 (*nekl-4p∷nekl-4∷gfp*), *nekl-4* plus a 1609bp upstream promoter region was amplified from genomic DNA using primers with homology to pPD95.75 containing *gfp.* The stop codon for *nekl-4* was removed using the reverse primer. pPD95.75 was amplified using primers with homology to *nekl-4.* Fragments were joined using Gibson assembly [95].

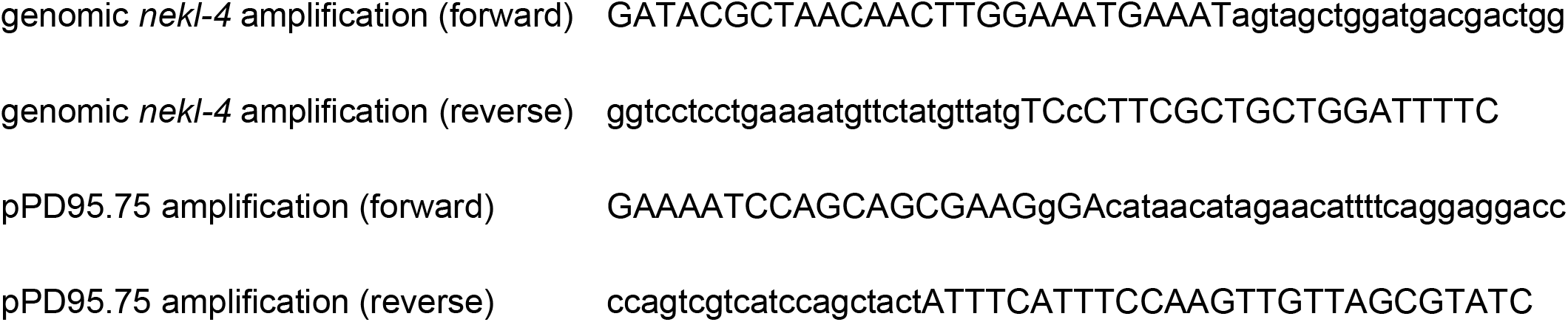

### CRISPR constructs

For creation of the *nekl-4∷mneongreen* and *nekl-4∷mscarlet-I* endogenous CRISPR tags, we followed the Mello Lab protocol [96] and used pdsDNA [97] that included a short flexible linker sequence between the 3’ end of *nekl-4* and the start of *mneongreen* or *mscarlet-I.* The guide sequence was designed using CRISPOR [98,99] and silent mutation sites for PAM modification and diagnosis were located using WatCut silent mutation scanning [100]. Reagents used are as follows. dg357 was a gift from Dominique A. Glauser and was used in accordance with the *C. elegans* group license with AlleleBiotech. pSEM90 was a gift from Thomas Boulin.

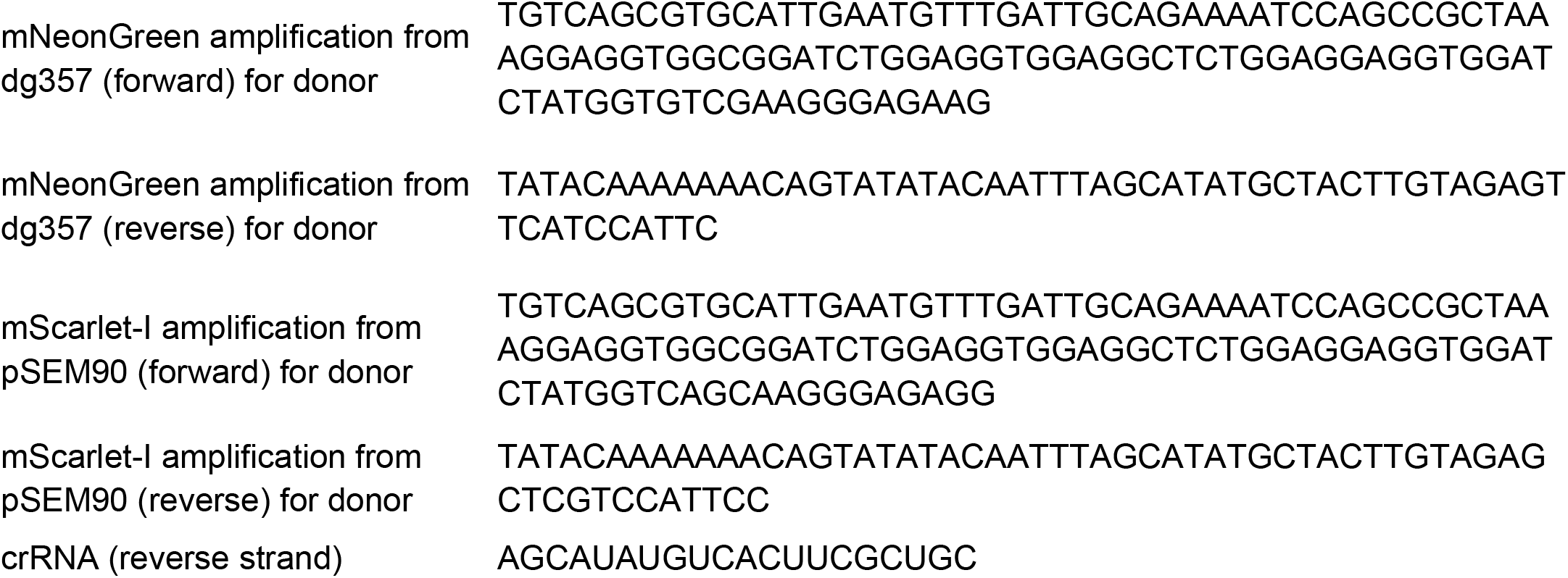

---

**Supplemental Table 1.**
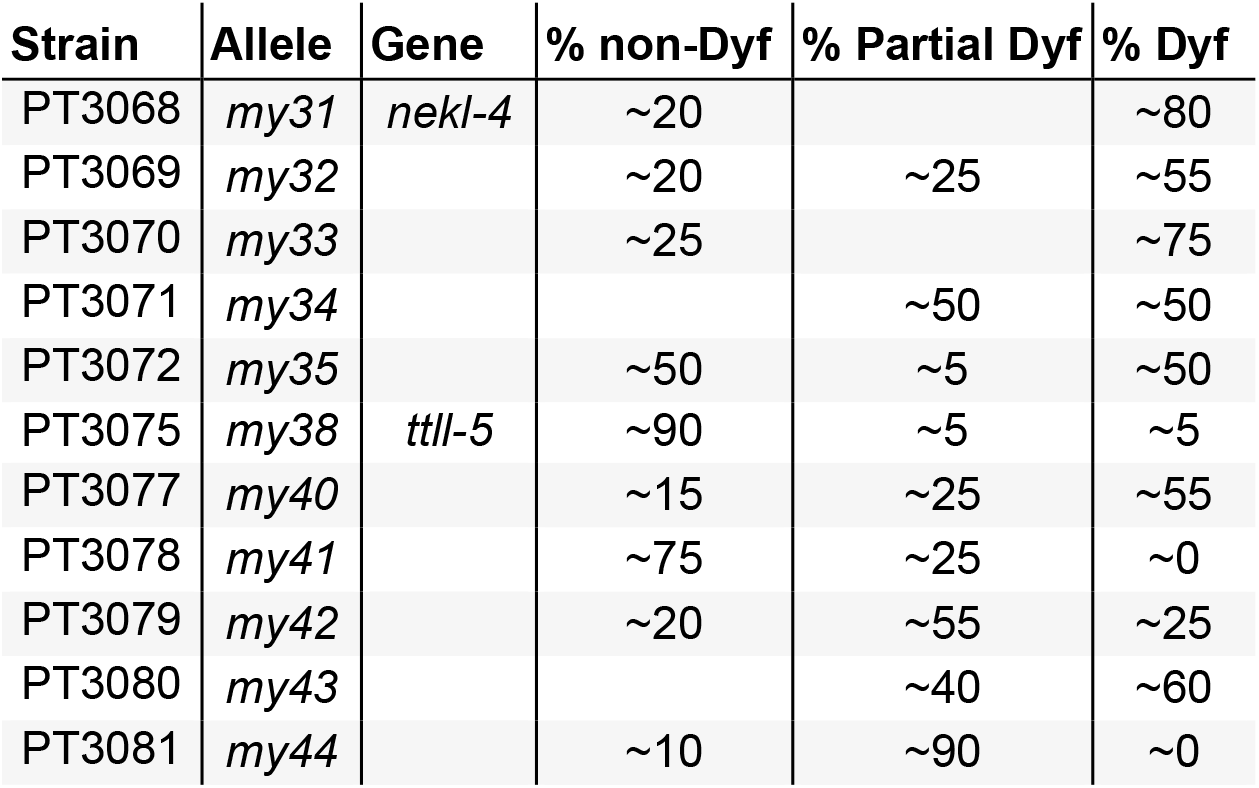
List of independent *ccpp-1* suppressor isolates. Strains contain the allele mentioned and the *ccpp-1(ok1821)* deletion. The approximate percentage of non Dyf, Partial Dyf, and Dyf animals of each strain is indicated. At least 20 animals were tested for Dye filling for each strain. Although a single P_0_ plate produced *my42* and *my43*, we considered them independent isolates because their phenotypes are different from one another.

**Supplemental Table 2.** Mutations examined in this study. GKO; Gene Knockout Consortium. CGC; Caenorhabditis Genetics Center. NBRP; National Bio-Resource Project.

| Gene | Allele | Molecular Lesion | Source |
| --- | --- | --- | --- |
| *ccpp-1* | *ok1821* | 590 AA in-frame deletion; 5' regulatory region and 6 exons | GKO |
| *ttll-4* | *tm3310* | 137 AA out-of-frame deletion, 2 AA insertion; TTL domain | NBRP |
| *ttll-5* | *my38* | L338F; TTL domain | this study |
|  | *tm3360* | 277 AA in-frame deletion, 4 AA insertion; TTL domain | NBRP |
| *ttll-11* | *tm4059* | 169 AA in-frame deletion, TTL domain | NBRP |
|  | *gk482* | 180 AA out-of-frame deletion; start and first exon of isoform b | CGC |
| *nekl-4* | *my31* | W199X; 5th exon | this study |
|  | *tm4910* | 135 AA in-frame deletion, replaced with Q; kinase domain | NBRP |

**Supplemental Movie 1.**
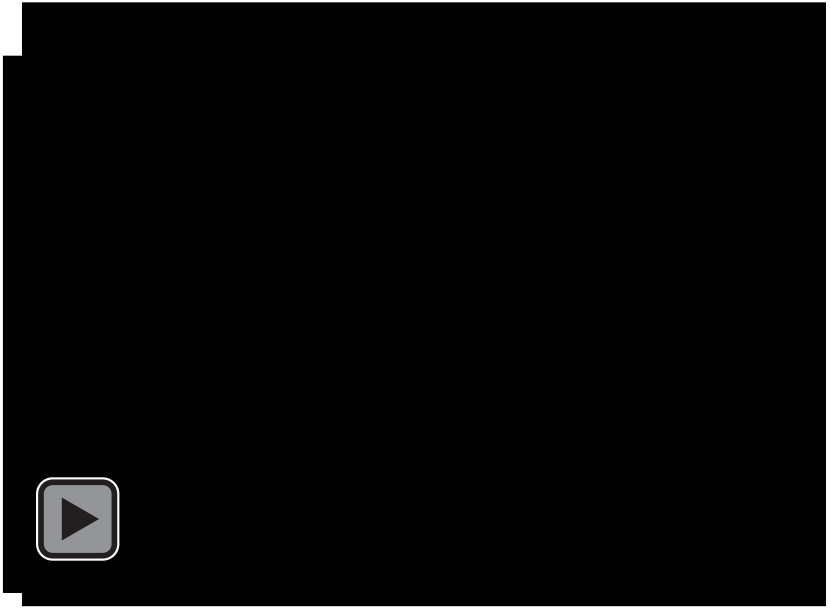
NEKL-4∷mNeonGreen movement in the amphid distal dendrites. Time lapse of a single z slice with widefield microscope. Images acquired at 4.34 fps and shown at 20 fps.

**Supplemental Figure 1.**
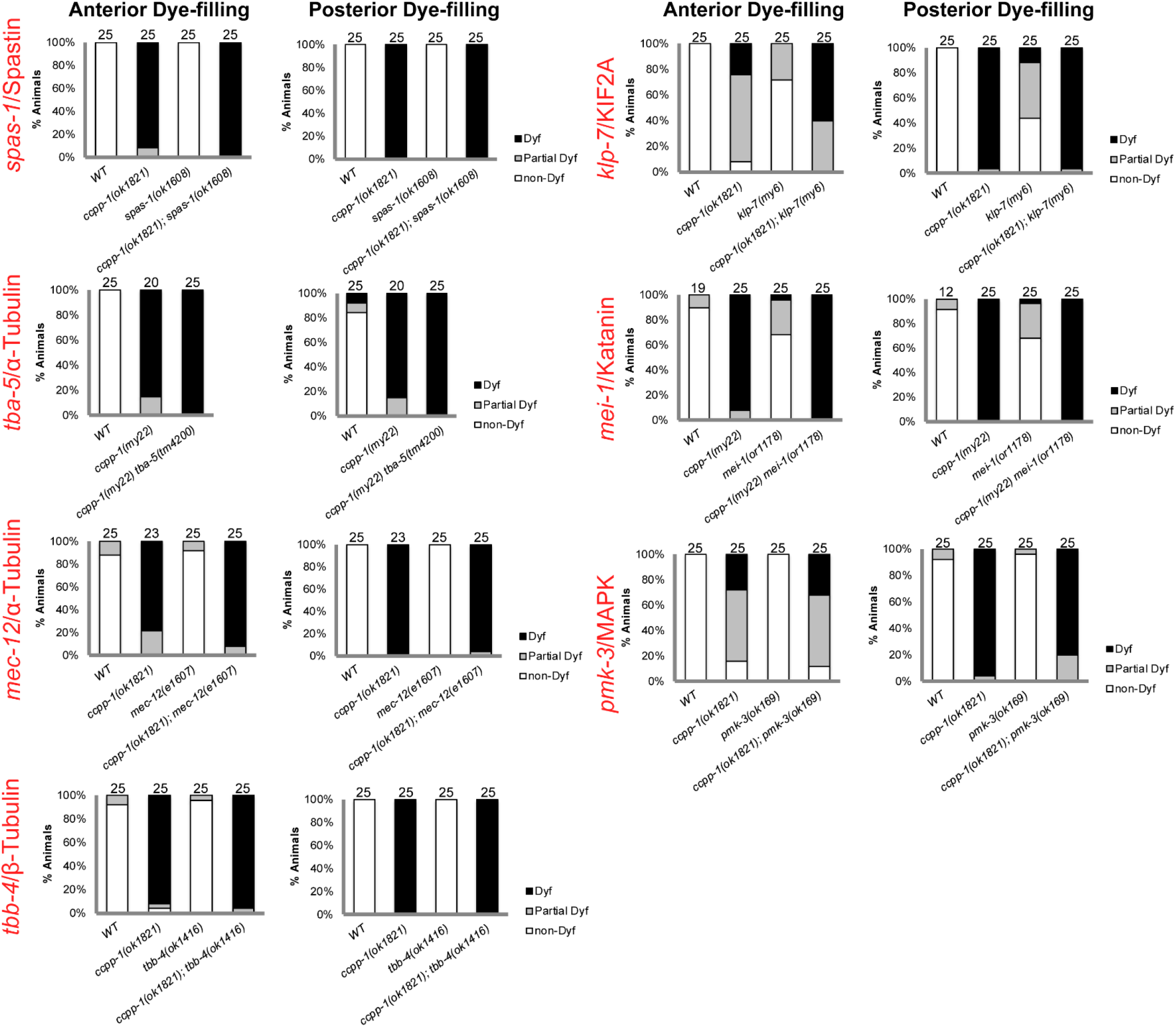
Candidate gene mutations that did not suppress the *ccpp-1* dye-filling defect. For experiments involving the *mei-1(or1178)* temperature-sensitive allele, eggs from plates kept at the permissive temperature (15°C; [101]) were picked to fresh plates and shifted to 25°C after hatching for 4 days before scoring.

**Supplemental Figure 2.**
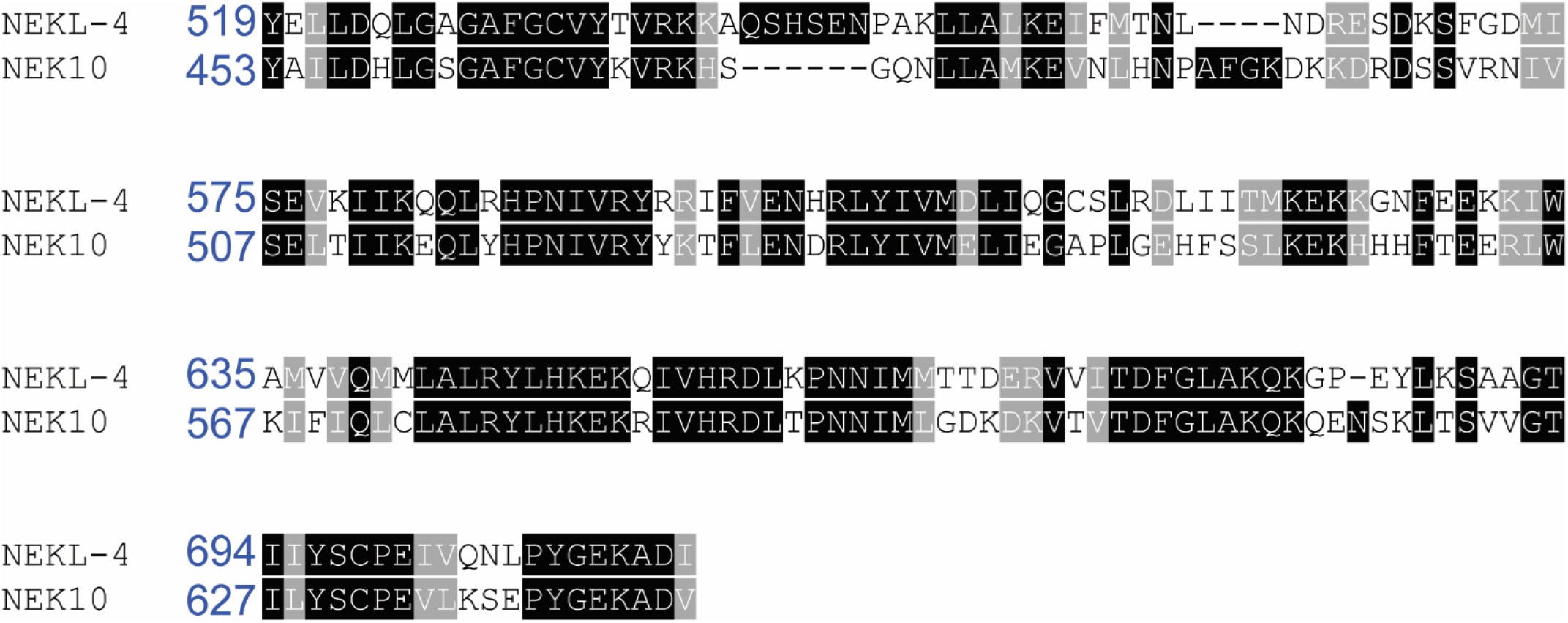
Alignment of *C. elegans* NEKL-4 and human NEK10 kinase domains.

**Supplemental Figure 3.**
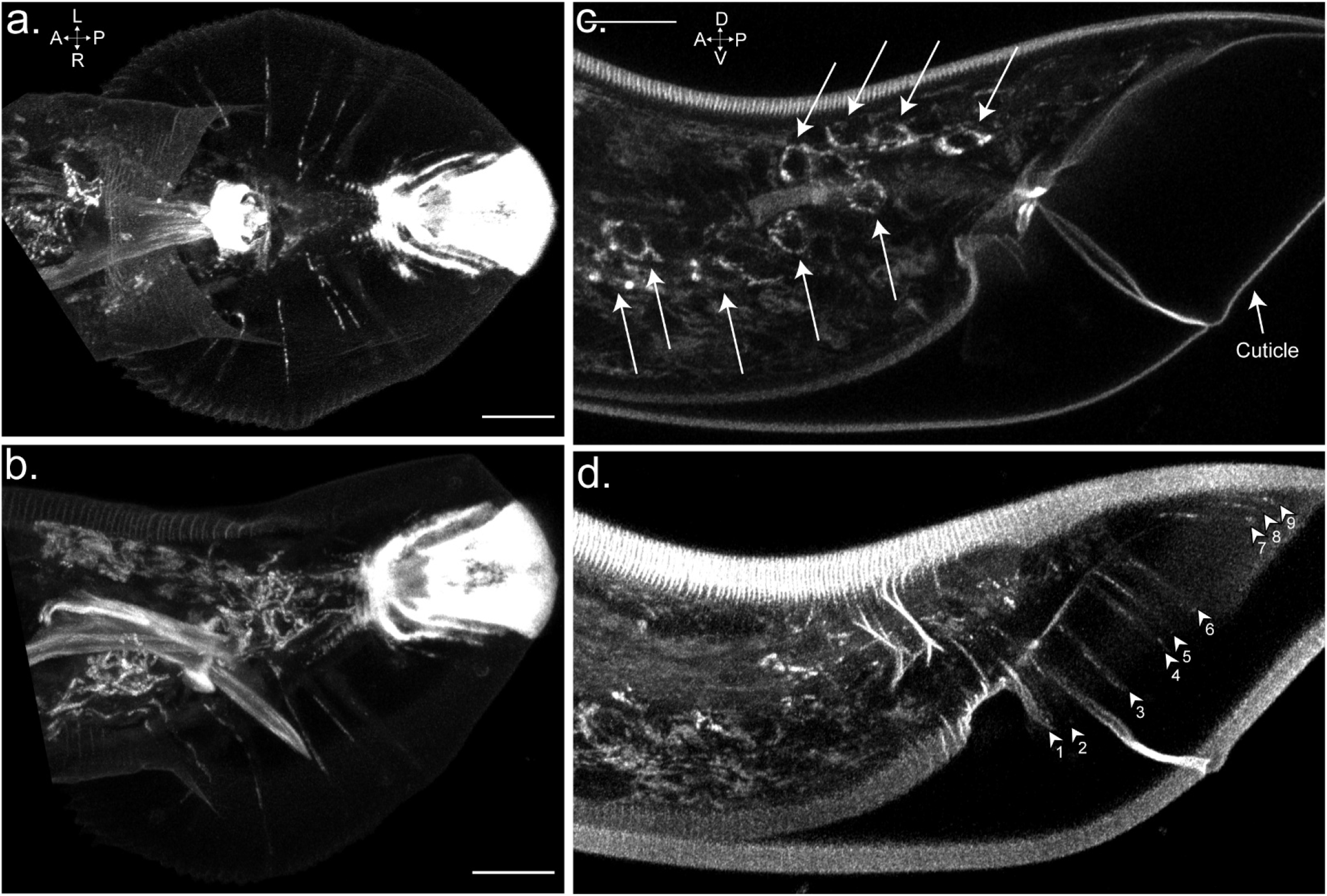
NEKL-4 is localized to specialized ciliated sensory neurons in the male tail. **a-b.** Localization of NEKL-4∷mNeonGreen in adult males. Tail is flat against the coverslip in **a.** to visualize ray dendrites. Tail in **b.** is folded but the filamentous pattern in ray cell bodies is visible. **c-d.** Localization of NEKL-4∷mNeonGreen in L4 molt males. Both images are different sections from the same z-stack. Arrows indicate ray cell bodies, arrowheads indicate ray dendrites (numbered). Scale = 10μm.

**Supplemental Table 3.** Summary of suppression of *ccpp-1* phenotypes by mutations examined in this study. + indicates any degree of suppression of the *ccpp-1* phenotype, - indicates no suppression of the *ccpp-1* phenotype, n/a indicates not tested.

| Suppressor gene | Allele | Dye-filling | GT335 staining | KAP-1 localization |
| --- | --- | --- | --- | --- |
| ***ttl-4*** | *tm3310* | + | + | n/a |
| ***ttl-5*** | *my38* | + | + | n/a |
|  | *tm3360* | + | + | + |
| ***ttl-11*** | *tm4059* | + | - | + |
|  | *gk482* | - | + | n/a |
| ***nekl-4*** | *my31* | + | + | n/a |
|  | *tm4910* | + | - | - |

**Supplemental Table 4.**
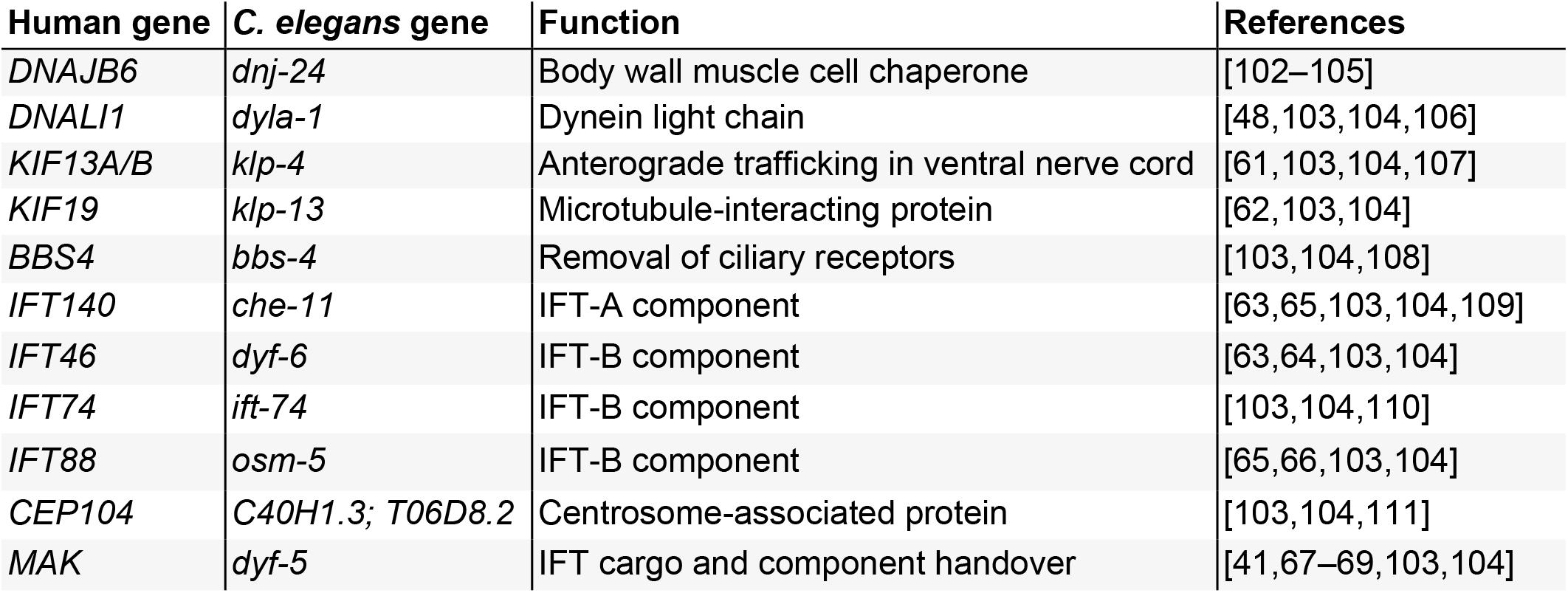
Potential NEKL-4 substrates. Based on phosphoproteomics data for NEK10 from Chivukula *et al*, Figure 4f [26]. Original list contains genes with phosphopeptides depleted more than twofold upon NEK10 deletion in human bronchial epithelial cell (HBEC) culture. This list contains genes with known *C. elegans* homologs from the following relevant classes in the NEK10 list: axonemal dyneins and assembly factors, kinesins, intraflagellar transport, ciliary length control.

| Human gene | *C. elegans* gene | Function | References |
| --- | --- | --- | --- |
| *DNAJB6* | *dnj-24* | Body wall muscle cell chaperone | [102–105] |
| *DNALI1* | *dyla-1* | Dynein light chain | [48,103,104,106] |
| *KIF13A/B* | *klp-4* | Anterograde trafficking in ventral nerve cord | [61,103,104,107] |
| *KIF19* | *klp-13* | Microtubule-interacting protein | [62,103,104] |
| *BBS4* | *bbs-4* | Removal of ciliary receptors | [103,104,108] |
| *IFT140* | *che-11* | IFT-A component | [63,65,103,104,109] |
| *IFT46* | *dyf-6* | IFT-B component | [63,64,103,104] |
| *IFT74* | *ift-74* | IFT-B component | [103,104,110] |
| *IFT88* | *osm-5* | IFT-B component | [65,66,103,104] |
| *CEP104* | *C40H1.3; T06D8.2* | Centrosome-associated protein | [103,104,111] |
| *MAK* | *dyf-5* | IFT cargo and component handover | [41,67–69,103,104] |

**Supplemental Table 5.** Strains used in this work. All strains were generated in the Barr laboratory except CB1490 (Hodgkin laboratory), ROH11 and ROH12 (O’Hagan laboratory).

| Strain | Genotype |
| --- | --- |
| CB1490 | *him-5(e1490)V* |
| PT443 | *myls1[pkd-2::gfp+unc-122p::gfp] pkd-2(sy606)IV;him-5(e1490)V* |
| PT3309 | *ccpp-1(ok1821)I;myls1[pkd-2::gfp+unc-122p::gfp] pkd-2(sy606)IV;him-5(e1490)V* |
| PT3310 | *ttl-4(tm3310)III;myls1[pkd-2::gfp+unc-122p::gfp] pkd-2(sy606)IV;him-5(e1490)V* |
| PT3311 | *ccpp-1(ok1821)I;ttl-4(tm3310)III;myls1[pkd-2::gfp+unc-122p::gfp] pkd-2(sy606)IV;him-5(e1490)V* |
| PT3308 | *ttl-5(my38)V* |
| PT3283 | *ccpp-1(ok1821)I;ttl-5(my38)V* |
| PT3312 | *myls1[pkd-2::gfp+unc-122p::gfp] pkd-2(sy606)IV;ttl-5 (tm3360) him-5(e1490)V* |
| PT3313 | *ccpp-1(ok1821)I;myls1[pkd-2::gfp+unc-122p::gfp] pkd-2(sy606)IV;ttl-5 (tm3360) him-5(e1490)V* |
| PT2498 | *myls1[pkd-2::gfp+unc-122p::gfp] pkd-2(sy606) ttl-11(tm4059)IV;him-5(e1490)V* |
| PT2497 | *ccpp-1(ok1821)I;myls1[pkd-2::gfp+unc-122p::gfp] pkd-2(sy606) ttl-11(tm4059)IV;him-5(e1490)V* |
| PT2988 | *myls1[pkd-2::gfp+unc-122p::gfp] pkd-2(sy606) ttl-11b(gk482)IV;him-5(e1490)V* |
| PT2958 | *ccpp-1(ok1821)I;myls1[pkd-2::gfp+unc-122p::gfp] pkd-2(sy606) ttl-11b(gk482)IV;him-5(e1490)V* |
| PT3268 | *nekl-4(tm4910)III;myls1[pkd-2::gfp+unc-122p::gfp] pkd-2(sy606)IV;him-5(e1490)V* |
| PT3476 | *nekl-4(my31)III;him-5(e1490)V* |
| PT3269 | *ccpp-1(ok1821)I;nekl-4(tm4910)III;myls1[pkd-2::gfp+unc-122p::gfp] pkd-2(sy606)IV;him-5(e1490)V* |
| PT3068 | *ccpp-1(ok1821)I;nekl-4(my31)III;myls1[pkd-2::gfp+unc-122p::gfp] pkd-2(sy606)IV;him-5(e1490)V* |
| PT3289 | *pha-1(e2123)III;him-5(e1490)V;myEx909[nekl-4p::nekl-4::gfp + pBX]* |
| PT3404 | *ccpp-1(ok1821)I;nekl-4(my31)III;myls1[pkd-2::gfp+unc-122p::gfp] pkd-2(sy606)IV;him-5(e1490)V;myEx927[nekl-4p::nekl-4::gfp + unc-122p::rfp]* |
| PT3406 | *nekl-4(my51[nekl-4::mneongreen])III;him-5(e1490)V* |
| PT3484 | *nekl-4(my69[nekl-4::mscarlet])III;him-5(e1490)V;yhEx149[nphp-1::cfp+che-13::yfp+pRF4]* |
| PT3366 | *him-5(e1490)V;Ex[kap-1::gfp+pRF4]* |
| PT2151 | *ccpp-1(ok1821)I;him-5(e1490)V;Ex[kap-1::gfp+pRF4]* |
| PT3023 | *ttl-4(tm3310)III;Ex[kap-1::gfp+pRF4]* |
| PT3024 | *ttl-11(tm4059)IV;Ex[kap-1::gfp+pRF4]* |
| PT3027 | *ccpp-1(ok1821)I;ttl-11(tm4059)IV;Ex[kap-1::gfp+pRF4]* |
| PT3372 | *ttl-5 (tm3360) him-5(e1490)V;Ex[kap-1::gfp+pRF4]* |
| PT3373 | *ccpp-1(ok1821)I;ttl-5 (tm3360) him-5(e1490)V;Ex[kap-1::gfp+pRF4]* |
| PT3388 | *nekl-4(tm4910)III;him-5(e1490)V;Ex[kap-1::gfp+pRF4]* |
| PT3389 | *ccpp-1(ok1821)I;nekl-4(tm4910)III;him-5(e1490)V;Ex[kap-1::gfp+pRF4]* |
| PT3407 | *osm-3(p802)IV;him-5(e1490)V;Ex[kap-1::gfp+pRF4]* |
| PT1974 | *osm-3(p802)IV;him-5(e1490)V* |
| PT3270 | *nekl-4(tm4910)III;him-5(e1490)V* |
| ROH11 | *ttl-4(tm3310)III;osm-3(p802)IV;him-5(e1490)V* |
| ROH12 | *osm-3(p802)IV;ttl-5(tm3360) him-5(e1490)V* |
| PT3320 | *osm-3(p802) ttl-11(tm4059)IV;him-5(e1490)V* |
| PT3347 | *nekl-4(tm4910)III;osm-3(p802)IV;him-5(e1490)V* |
| PT3262 | *Ex[osm-3::gfp+pRF4]* |
| PT3252 | *ccpp-1(ok1821)I;Ex[osm-3::gfp+pRF4]* |
| PT3037 | *ccpp-1(ok1821)I;HW* |
| PT3075 | *ccpp-1(ok1821)I;ttl-5(my38)V* |

**Supplemental Table 6.** Strains used in supplemental material. All strains were generated in the Barr laboratory except BS3383, which was obtained from the Caenorhabditis Genetics Center (CGC).

| Strain | Genotype |
| --- | --- |
| PT3005 | *myls1[pkd-2::gfp+unc-122p::gfp] pkd-2(sy606)IV;spas-1(ok1608)V* |
| PT3006 | *ccpp-1(ok1821)I;myls1[pkd-2::gfp+unc-122p::gfp] pkd-2(sy606)IV;spas-1(ok1608)V* |
| PT3008 | *dpy-5(e61) ccpp-1(my22) tba-5(tm4200)I;myls1[pkd-2::gfp+unc-122p::gfp] pkd-2(sy606)IV;him-5(e1490)V* |
| PT2954 | *myls1[pkd-2::gfp+unc-122p::gfp] pkd-2(sy606)IV;him-5(e1490)V;tbb-4(ok1416)X* |
| PT2955 | *ccpp-1(ok1821)I;myls1[pkd-2::gfp+unc-122p::gfp] pkd-2(sy606)IV;him-5(e1490)V;tbb-4(ok1416)X* |
| PT3144 | *mec-12(e1607)III;myls1[pkd-2::gfp+unc-122p::gfp] pkd-2(sy606)IV;him-5(e1490)V* |
| PT3143 | *ccpp-1(ok1821)I;mec-12(e1607)III;myls1[pkd-2::gfp+unc-122p::gfp] pkd-2(sy606)IV;him-5(e1490)V* |
| BS3383 | *pmk-3(ok169)IV* |
| PT2673 | *klp-7(my6)III;myls1[pkd-2::gfp+unc-122p::gfp] pkd-2(sy606)IV;him-5(e1490)V* |
| PT2672 | *ccpp-1(ok1821)I;klp-7(my6)III;myls1[pkd-2::gfp+unc-122p::gfp] pkd-2(sy606)IV;him-5(e1490)V* |
| PT3014 | *dpy-5(e61) ccpp-1(my22) mei-1(or1178)I;myls1[pkd-2::gfp+unc-122p::gfp] pkd-2(sy606)IV;him-5(e1490)V* |

## Author contributions

K. M. P.; M. M. B.; and R. O. conceived the study and wrote the manuscript. J. S. A. wrote the manuscript. K. M. P.; A. G.; J. D. W.; S. B.; M. M.; W. Z.; A. G.; H. S.; and R. O. performed experiments and analyzed data. N. R. constructed strains and analyzed data. J. D. W.; A.G.; H. S.; M. M. B.; and R. O. analyzed data and provided funding.

## Acknowledgements

This work was funded by NJCSCR CSCR15IRG014 to R. O.; NIH DK116606 to M. M. B.; Institutional Research and Academic Career Development Award (IRACDA) 1K12GM093854 to J. D. W; and the Intramural Research Program of the National Institutes of Health, National Institute of Diabetes and Digestive and Kidney Diseases (NIDDK) (H.S. and A.G.). We thank members of the Barr lab, the Rutgers University *C. elegans* community, and K.P.’s thesis committee Marc Gartenberg, Monica Driscoll, and Bonnie Firestein for advice and helpful feedback; Antonina Roll-Mecak for insight on TTLL-5 structure; and Helen Ushakov and Gloria Androwski for technical assistance. We thank Dominique A. Glauser and Thomas Boulin for plasmids. We also thank WormBase (U41 HG002223) and WormAtlas (R24 OD010943) for online resources. Some strains were provided by the CGC, which is funded by NIH Office of Research Infrastructure Programs (P40 OD010440), or by the National Bio-Resource Project of the MEXT, Japan.

